# Oncogenic Gene Fusions in Non-Neoplastic Precursors as Evidence that Bacterial Infection Initiates Prostate Cancer

**DOI:** 10.1101/2020.07.27.224154

**Authors:** Eva Shrestha, Jonathan B. Coulter, William Guzman, Busra Ozbek, Luke Mummert, Sarah E. Ernst, Janielle P. Maynard, Alan K. Meeker, Christopher M. Heaphy, Michael C. Haffner, Angelo M. De Marzo, Karen S. Sfanos

**Affiliations:** Department of Pathology, Johns Hopkins University School of Medicine, Baltimore, MD 21287; Sidney Kimmel Comprehensive Cancer Center, Johns Hopkins University School of Medicine, Baltimore, MD 21287; Department of Art as Applied to Medicine, Johns Hopkins University School of Medicine, Baltimore, MD 21287; Department of Urology, James Buchanan Brady Urological Institute, Johns Hopkins University School of Medicine, Baltimore, MD 21287

**Author notes:** Correspondence to: Karen S. Sfanos, Department of Pathology, Johns Hopkins University School of Medicine, 1550 Orleans Street, CRB II Rm. 143, Baltimore, MD 21287. Boston University School of Medicine, Boston, MA. Division of Human Biology, Fred Hutchinson Cancer Research Center, Seattle, WA.

## Abstract

Prostate adenocarcinoma is the second most commonly diagnosed cancer in men worldwide and the initiating factors are unknown. Oncogenic TMPRSS2:ERG (ERG+) gene fusions are facilitated by DNA breaks and occur in up to 50% of prostate cancers^1,2^. Infection-driven inflammation is implicated in the formation of ERG+ fusions^3^, and we hypothesized that these fusions initiate in early inflammation-associated prostate cancer precursor lesions, such as proliferative inflammatory atrophy (PIA), prior to cancer development. We investigated whether bacterial prostatitis is associated with ERG+ precancerous lesions in unique cases with active bacterial infections at time of radical prostatectomy. We identified a high frequency of ERG+ non-neoplastic-appearing glands in these cases, including ERG+ PIA transitioning to early invasive cancer. We verified TMPRSS2:ERG genomic rearrangements in precursor lesions using tri-color fluorescence *in situ* hybridization. Identification of rearrangement patterns combined with whole prostate mapping in 3 dimensions confirmed multiple (up to 8) distinct ERG+ precancerous lesions in infected cases. Finally, we identified the pathogen-derived genotoxin colibactin as a potential source of DNA breaks in clinical cases as well as cultured prostate cells. Overall, we provide evidence that bacterial infections initiate driver gene alterations in prostate cancer. Furthermore, infection-induced ERG+ fusions are an early alteration in the carcinogenic process and PIA may serve as a direct precursor to prostate cancer.

---

TMPRSS2:ERG gene fusions are an early prostate cancer oncogenic event, as essentially all invasive cancer cells within an ERG positive (ERG+) cancer have the same cytogenetically defined rearrangement^4^. Likewise, in mouse models, transgenic expression of ERG in luminal prostate epithelial cells induces prostatic intraepithelial neoplasia (PIN)^5,6^, and carcinoma when combined with other genomic alterations^7-9^. Inflammation induced by bacterial lipopolysaccharide (LPS) has been demonstrated in both *in vitro* and *in vivo* models to induce the formation of TMPRSS2:ERG gene fusions^3^. Prostatic inflammation may likewise drive the formation of proliferative inflammatory atrophy (PIA), a putative prostate cancer precursor lesion^10,11^. In addition to stimulating inflammation, bacteria can produce potent genotoxins that incite DNA damage, as has been described in the pathogenesis of colon cancer^12-18^. Therefore, we questioned whether bacterial infections are associated with oncogenic TMPRSS2:ERG gene fusions in human prostate specimens. Furthermore, we hypothesized that infection-induced TMPRSS2:ERG fusions may initiate in early inflammation-associated prostate cancer risk factor lesions, such as PIA, prior to cancer development.

We began by identifying a series of radical prostatectomy specimens that were suspicious for prostatic infections at the time of surgery. Our rationale was that even though these specimens had pre-existing cancer, we could examine the non-neoplastic regions of these highly inflamed cases to investigate the effects of infection-induced inflammation on the human prostate. Florid acute or granulomatous chronic inflammation observed at radical prostatectomy is rare, and likely indicative of an active prostate infection. In screening 1,341 cases (see Methods), we identified 15 cases that were suspicious for the presence of an active prostate infection at the time when the radical prostatectomy was performed (Extended Data Fig. 1 and Extended Data Table 1). We performed 100% prostate sampling and mapping of all invasive carcinoma foci in 3 dimensions (3D) in these cases.

**Fig. 1.**
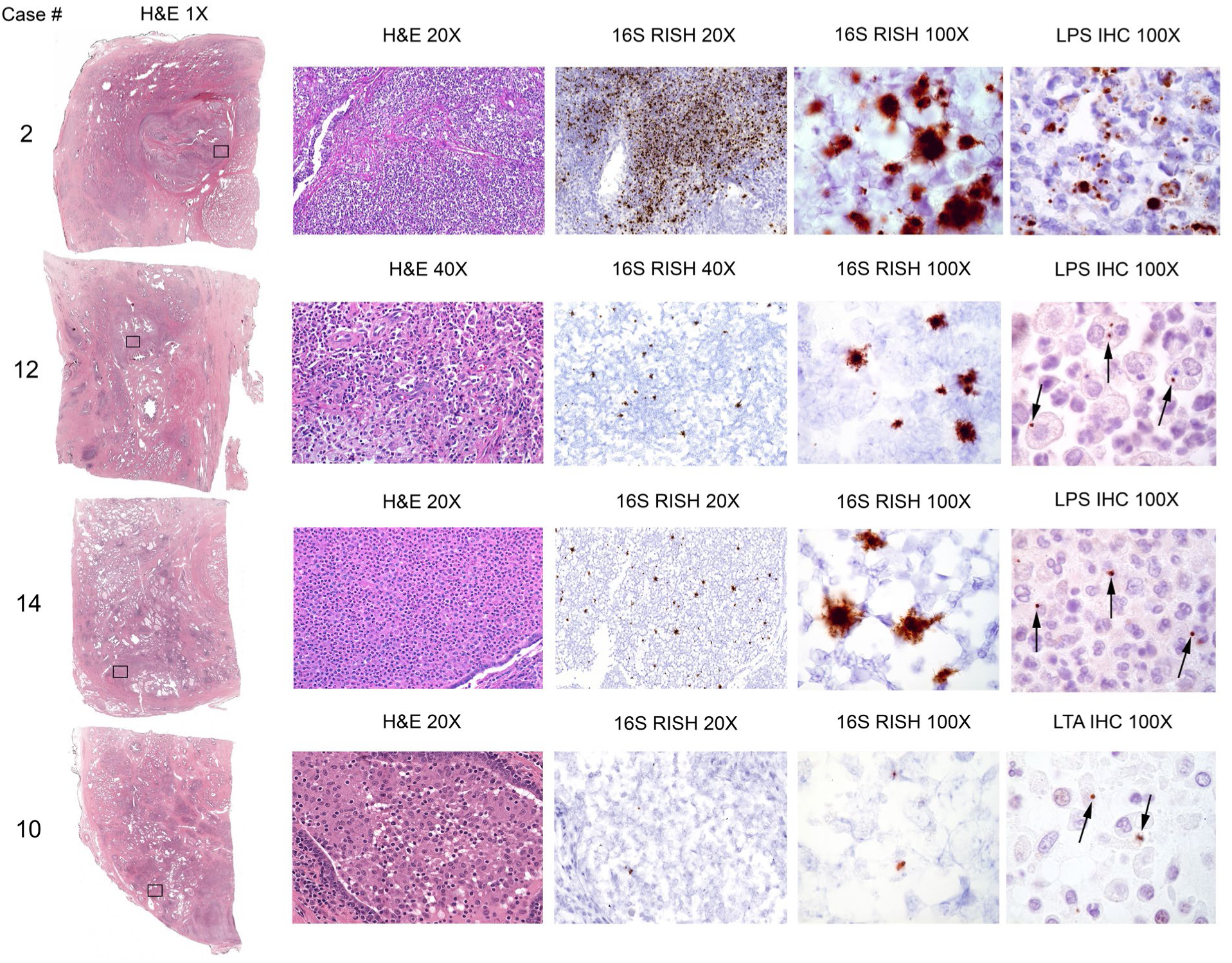
Detection of bacteria in highly inflamed radical prostatectomy specimens. Examples of detection of bacteria by 16S RISH (universal detection of bacteria), LPS IHC (specific for Gram negative bacteria, case 2, 12, and 14) and LTA IHC (specific for Gram positive bacteria, case 10). Bacterial cells were often observed within infiltrating immune cells (arrows). Objective magnification denoted.

To further evaluate whether these cases harbored bacterial infections, we assessed all blocks that contained acute inflammation (defined as the presence of neutrophils, average 9 blocks per case) using RNA *in situ* hybridization (RISH) with universal probes for bacterial 16S ribosomal RNA (rRNA) as well as immunohistochemistry (IHC) for LPS (Gram negative bacteria) and lipoteichoic acid (LTA, Gram positive bacteria). RISH and IHC assay validation data are in Extended Data Fig. 2. We detected bacteria in seven of the cases, six of which were LPS positive and one that was LTA positive (Fig. 1, Extended Data Table 1, and Extended Data Fig. 3). Importantly, even though bacteria were not identified in the remaining cases, the cases are still highly suspicious for infection due to the presence of florid inflammation. It is possible that the infectious agent cleared prior to the radical prostatectomy, since in the positive cases only a small number of glands were bacteria-positive, despite florid inflammation across the bacteria-negative glands. Special stains for acid fast bacilli (auramine/rhodamine) and fungi (methenamine silver) were performed on a subset of the cases during the initial diagnostic workup and were negative (Extended Data Table 1). Non-inflamed regions of normal appearing prostate and prostate adenocarcinoma were all negative for bacteria (Extended Data Fig. 4). When present, bacteria were largely confined to glandular lumens where neutrophils and macrophages were present, and were often observed to be internalized by these cells (Fig. 1, Extended Data Fig. 3).

**Fig. 2.**
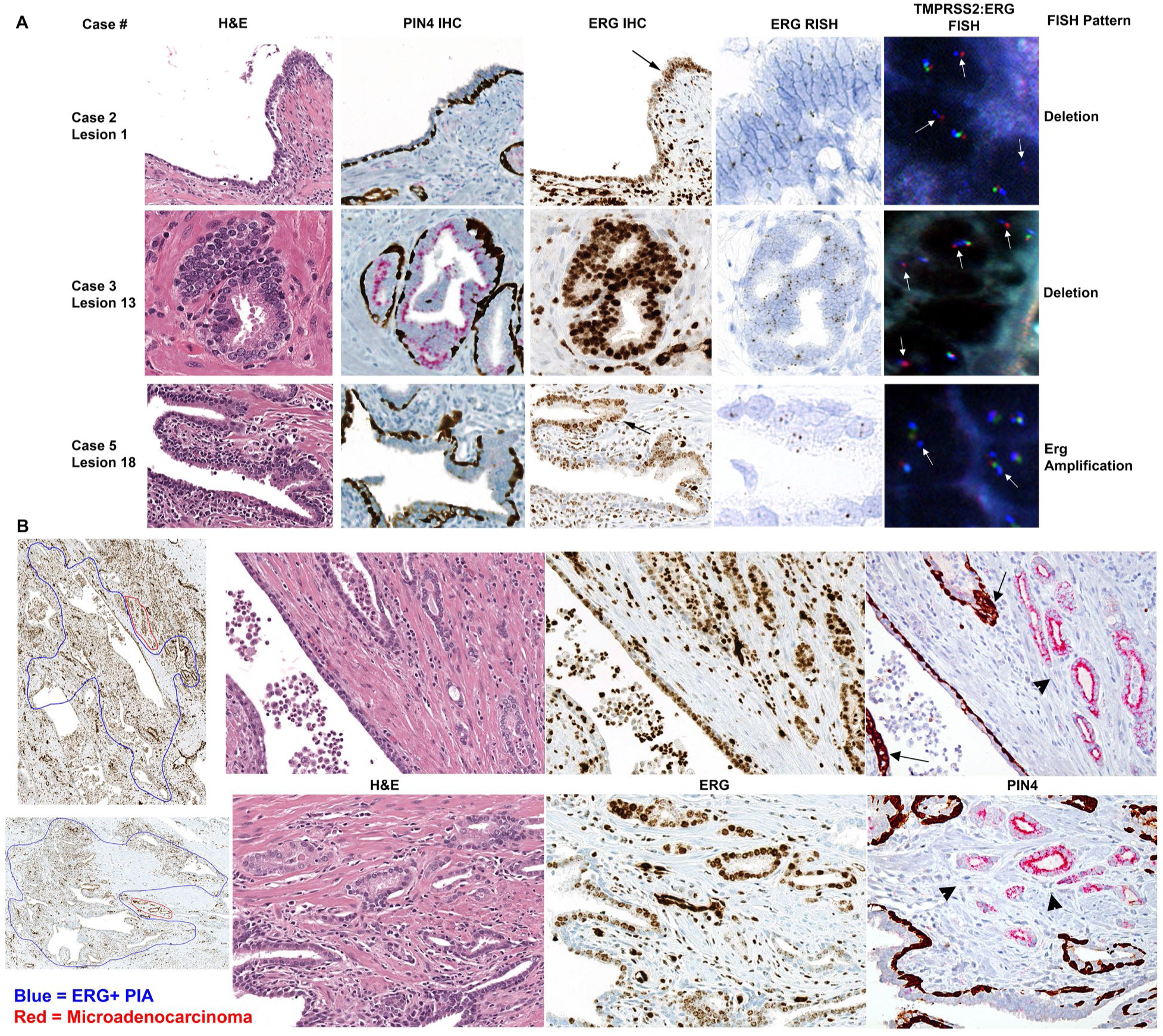
ERG expression from TMPRSS2:ERG gene fusions present in luminal epithelial cells in PIA (lesion 1 and 18) and LGPIN (lesion 13) in highly inflamed radical prostatectomy specimens. (A) Representative images from case 2, 3, and 5 showing intact basal cells indicative of a non-neoplastic gland (PIN4 IHC) and positive ERG IHC, RISH, and FISH in luminal epithelial cells (200X magnification). Black arrows point to magnified (400X magnification) regions in ERG RISH. White arrows point to TMPRSS2:ERG genomic rearrangements detected via FISH as denoted (400X magnification). Red probe is located in the distal TMPRSS2 gene region, green probe is located in the proximal TMPRSS2 gene region, and blue probe is located in the ERG (21q22) gene region. (B) Large regions of ERG+ PIA (circled in blue in first panel of ERG IHC, 40X magnification) with small regions of adjacent microadenocarcinoma (circled in red) from case 2 (top row) and case 5 (bottom row). Infiltrating immune cells and endothelial cells are also ERG+. H&E, ERG IHC, and PIN4 IHC (all 200X magnification) demonstrate ERG+ PIA next to ERG+ microadenocarcinoma (arrowheads). Arrows denote intermediate cells in PIN4 IHC.

**Fig. 3.**
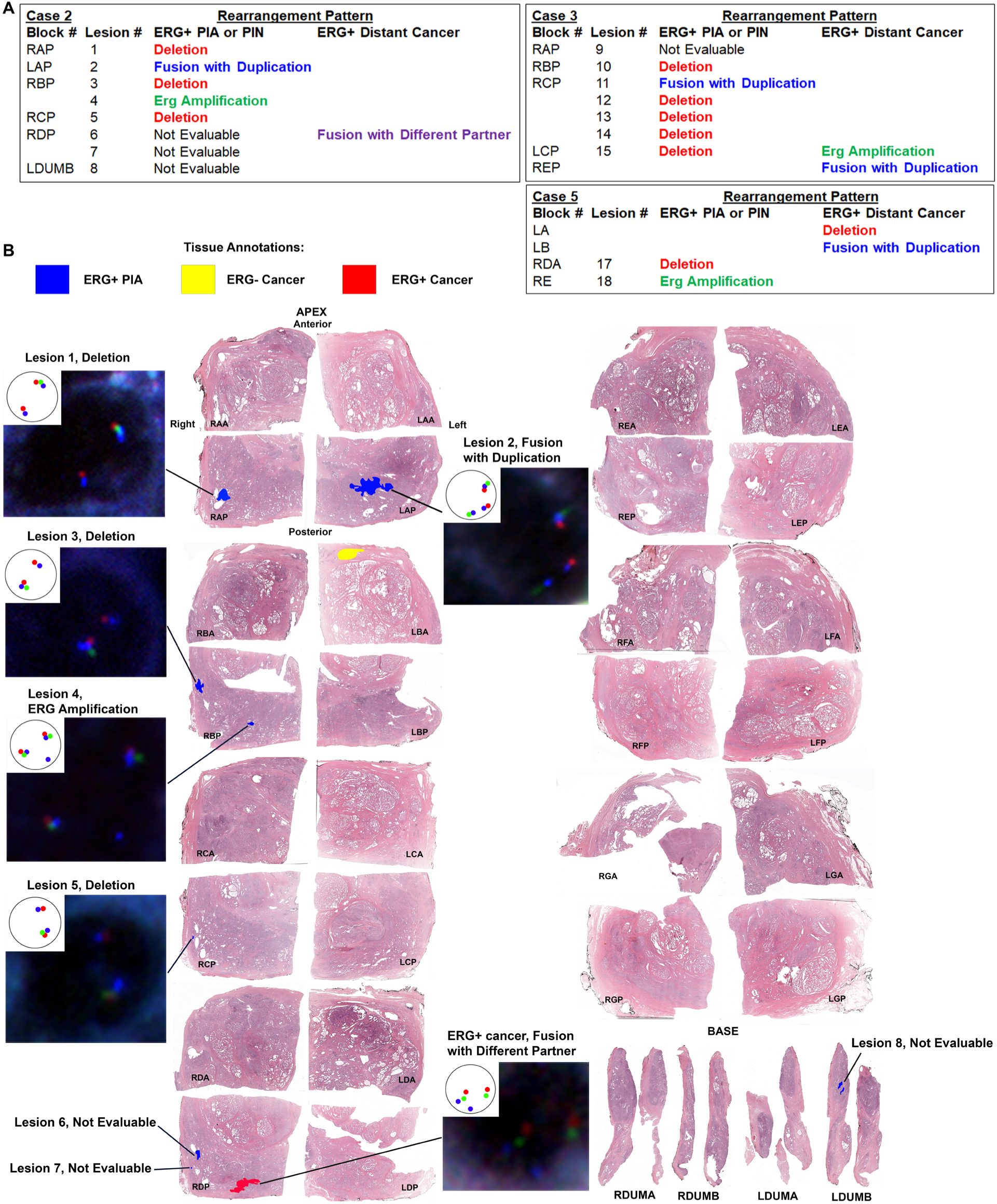
Evaluation and spatial mapping of TMPRSS2:ERG genomic rearrangement patterns in separate PIA and PIN foci versus distant ERG+ cancer. (A) Results summary of TMPRSS2:ERG FISH analysis on each ERG+ lesion found in case 2, 3, and 5. Note that ERG+ PIA and LGPIN/HGPIN foci are often a different fusion pattern than the distant ERG+ cancer in the case, even if on the same block. Adjacent ERG-benign glands to ERG+ glands were assessed for TMPRSS2:ERG fusions in an identical manner to the ERG+ glands. All ERG-benign glands assessed did not contain TMPRSS2:ERG rearrangements. (B) Spatial mapping of TMPRSS2:ERG rearrangement patterns in case 2. Pattern shown in example nuclei is depicted in the inset. Red probe is located in the distal TMPRSS2 gene region, green probe is located in the proximal TMPRSS2 gene region, and blue probe is located in the ERG (21q22) gene region.

**Fig. 4.**
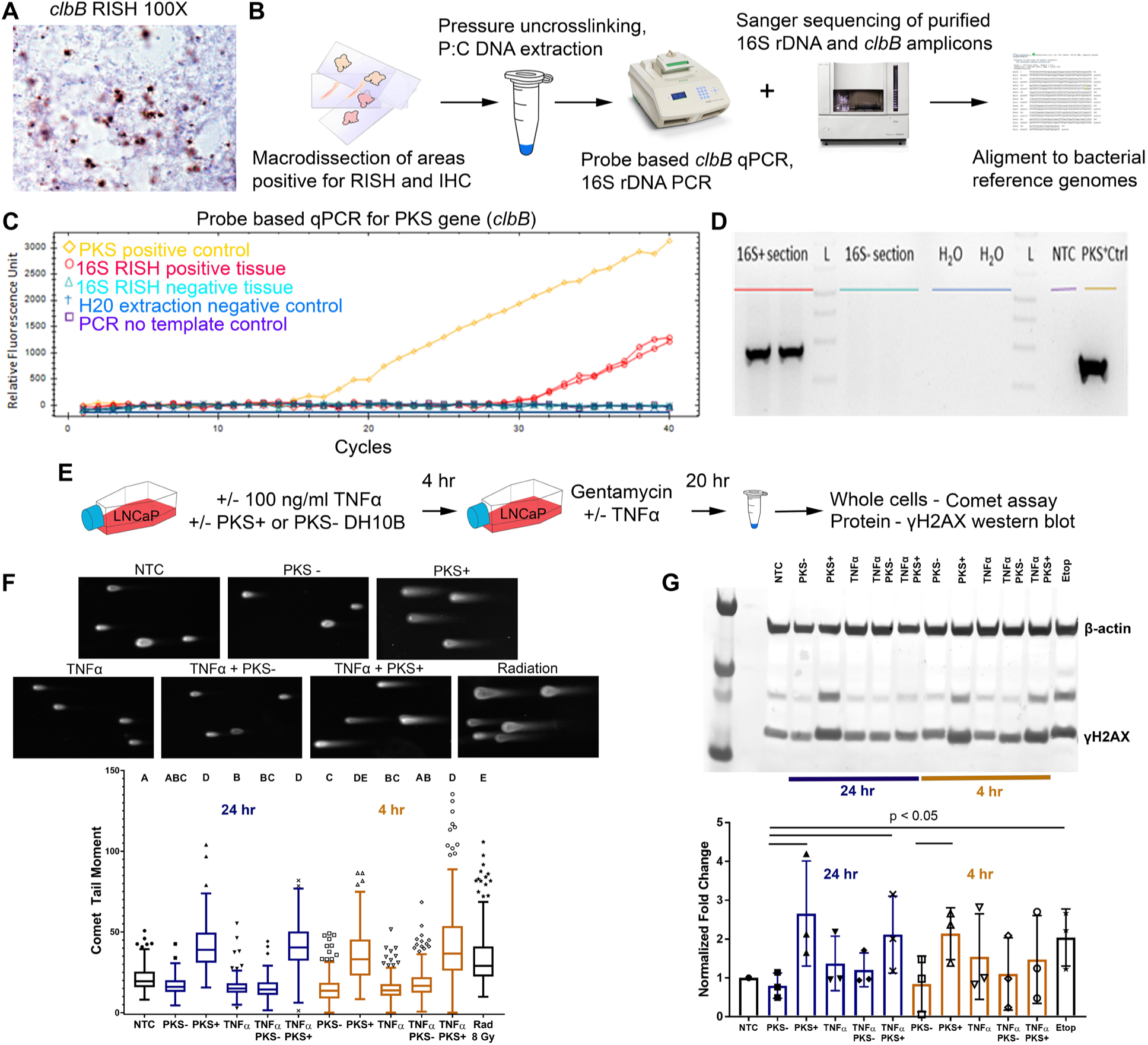
Exposure of prostate cancer cells to colibactin (PKS) induces DNA double strand breaks (DSB) as assessed by comet assay and western blot for γH2AX. (A) Example of PKS expression in the bacteria in case 2 (1,000 X magnification) as visualized by *cblB* RISH. (B) Strategy for bacterial DNA isolation from FFPE tissues, 16S rDNA sequencing, and PKS quantitative PCR (qPCR). (C) Detection of colibactin *clbB* gene by qPCR in DNA extracted from a macrodissected 16S RISH positive tissue area from case 2. (D) Agarose gel image of PCR products from PKS qPCR. (E) Experimental outline for *in vitro* colibactin exposure experiments. (F) Top, representative images of comet moments for each experimental condition. Comet moments are visibly induced in LNCaP cells by 8 Gy ionizing radiation exposure or 4 hr infection with PKS+ *E*. *coli*. NTC = no treatment control. Bottom, quantification of comet tail moments after exposure of LNCaP cells to PKS+ *E*. *coli* induces DNA DSB at or greater than exposure to 8 Gy radiation 4 hr and 24 hr post-exposure. The addition of TNFα does not increase DSB in this model system. Representative results of the experiment are shown. One way non-parametric ANOVA with Kruskal-Wallis multiple comparison test was performed. Differing letters denote statistical significance (p < 0.05). The statistical significance of PKS+ *E*. *coli*-treated groups at 4 hours post-infection was variable across three biological replicates. The trend at 24 hours was consistent across four biological replicates. (G) Top, western blot for γH2AX likewise demonstrates a > 2-fold increase in DSB repair at 4 hr and 24 hr after PKS+ *E*. *coli* exposure. Etop = etoposide (positive control). Bottom, densitometry quantification of western blot images. The results are normalized to β-actin levels and no treatment control (NTC) and are representative of three biological replicates. Statistically significant differences noted (p < 0.05, one-tailed t-test).

Next, we used these cases to question whether bacterial prostatitis is associated with the presence of TMPRSS2:ERG gene fusions in early precursor lesions. Isolated ERG+ high-grade PIN (HGPIN) occurs at a rare frequency between 2% in prostatectomies^19^ and 7% in cystoprostatectomies^20^. ERG+ low-grade PIN (LGPIN) or benign acini are likewise exceptionally rare^19,21^. ERG+ PIA has not been previously described^22^. We performed ERG IHC (Extended Data Fig. 6) on 110 whole tissue sections from 64 unselected (not selected based on amount of inflammation) radical prostatectomy specimens and found isolated ERG+ LGPIN in one case, ERG+ PIA in one case, and both ERG+ LGPIN and PIA in one case (3 cases total, 4.7%, Extended Data Fig. 7). We define “isolated” ERG+ PIA or LGPIN as glands found on blocks (as well as the adjacent blocks) with no ERG+ cancer or > 9 mm away from ERG+ cancer, an important consideration in establishing that the ERG+ cells in the PIA or PIN foci are not representative of retrograde invasion of cancer cells into benign acini^23^. Our 3D mapping was used to avoid this possibility.

In stark contrast to the studies reported by Young *et al*.^19^, Furusato *et al*.^21^, and our assessment of whole tissue sections from unselected cases, we identified isolated ERG+ luminal epithelial cells in PIA and/or LGPIN in 7 (46.7%) of the 15 mapped cases in the highly inflamed/infection cohort (p < 0.0001 compared to unselected cases, chi-square test, Fig. 2A and Extended Data Fig. 8) when we examined ERG IHC on 126 whole tissues sections. We subsequently performed ERG IHC on all remaining blocks from the seven cases where we found ERG+ PIA and/or LGPIN to determine the spatial orientation of the ERG+ PIA or LGPIN foci to ERG+ or ERG-cancer in 3D (Extended Data Figs. 9-15). We identified multiple (up to 8) spatially distant (> 2 mm apart) foci of ERG+ PIA and/or LGPIN in 5 of the 7 cases (Extended Data Table 2). A total of 4 foci of ERG+ isolated HGPIN, 9 foci of isolated ERG+ LGPIN, and 15 foci of isolated ERG+ PIA were identified (Extended Data Table 2). These data strongly suggest infection-induced TMPRSS2:ERG fusions in PIA and LGPIN foci due to the high frequency and multiplicity of ERG+ foci found in these infected cases, which has not previously been described.

A dual IHC stain for Alpha-Methylacyl-CoA Racemase (AMACR) combined with high molecular weight cytokeratin (PIN4 IHC) confirmed the presence of basal cells in the ERG+ PIA or LGPIN foci (Fig. 2A), further indicating that these ERG+ foci are not cancer. In three of the cases, ERG+ PIA was identified in the absence of any ERG+ cancer in the blocks from the prostatectomy (Extended Data Fig. 11, 13, 15). *ERG* mRNA expression was also observed using RNA *in situ* hybridization (RISH, Extended Data Fig. 6) in the ERG+ PIA and LGPIN foci (Fig. 2A, Extended Data Fig. 16). We also confirmed ERG staining on a subset of the glands using IHC with a separate ERG antibody that targets a different epitope (C-versus N-terminus of ERG, Extended Data Fig. 17).

We noted with interest 7 foci in cases 2, 3, 4, and 5 where ERG+ PIA or LGPIN was directly adjacent to small clusters of glands that lacked basal cells and were apparently in the process of initial invasion (Fig. 2B, Extended Data Fig. 18) consistent with early invasive carcinoma (also called “microadenocarcinoma” ^24^). There was no other ERG+ cancer on the blocks with these lesions, or on either adjacent block in 3D. Furthermore, PIN4 IHC indicated that the ERG+ cells in the PIA lesions were AMACR negative, but the ERG+ cells in the budding adenocarcinoma were AMACR positive (Fig. 2B). These results suggest that the fusion event occurred in the PIA lesion and that TMPRSS2:ERG fusion events represent a very early, and perhaps even initiating event in prostate carcinogenesis, although requiring the acquisition of further oncogenic events to progress to adenocarcinoma. Intriguingly, we observed the presence of intermediate cells (luminal cells expressing the typical basal cell cytokeratins) in the PIA lesions that were adjacent to early invasive carcinoma (Fig. 2B, Extended Data Fig. 18). Intermediate cells are purported tumor initiating cells in prostate cancer^25,26^.

Since it is possible that ERG protein expression could be upregulated by mechanisms that do not involve fusion events, we verified that TMPRSS2:ERG genomic rearrangements were present in the ERG+ PIA and LGPIN glands. We used a triple color fluorescence *in situ* hybridization (FISH) assay designed to detect multiple scenarios of genomic alteration between the *TMPRSS2* and *ERG* loci on 21q22 (Extended Data Fig. 19) in three of the cases that had multiple ERG+ PIA and/or LGPIN foci (cases 2, 3, and 5). Non-neoplastic-appearing luminal epithelial cells that were positive by ERG IHC and *ERG* RISH were likewise positive for genomic alterations in the *TMPRSS2* and *ERG* loci (Fig. 2A, 3). Each of these cases had small ERG+ cancer that was distant to any of the ERG+ PIA or LGPIN foci (Extended Data Fig. 9, 10, and 12). In all three cases, the ERG+ cancer, in addition to being spatially distant to ERG+ PIA and LGPIN, contained a different rearrangement pattern than most or all of the ERG+ PIA and LGPIN foci in that same case (Fig. 3A,B). This finding further verifies that all or at least a subset the ERG+ PIA and LGPIN foci were distinct from any pre-existing ERG+ cancer present in a given case.

Two of the cases (case 2 and case 12) had bacteria present in large enough regions that we could macrodissect the tissue, extract DNA, and further identify the infecting species. Sequencing of a partial region of the 16S rRNA gene from these cases identified a bacterium in the family *Enterobacteriaceae*, which would also be consistent with the LPS IHC results (Fig. 1). Colibactin is a cryptic bacterial genotoxin produced by a non-ribosomal peptide synthetase and polyketide synthase gene cluster (*pks* island) carried in some strains of *E*. *coli* and closely related species^12^. Infection of human cells with colibactin-producing *E*. *coli* strains induces DNA damage including DNA adducts, DNA double-strand breaks, DNA cross-links, and genomic instability^12-16^. Colibactin-producing bacteria are present in the human gastrointestinal microbiota and associated with colorectal cancer^12,13,17,18,27^. Colibactin-producing *E*. *coli* are also known uropathogens and are common among isolates from men with bacterial prostatitis^28^. RISH for the colibactin *clbB* gene indicated that the infecting bacteria in both case 2 and case 12 were colibactin producers (Fig. 4A). The presence of the *pks* island in bacteria from these cases was further confirmed by probe-based quantitative PCR and sequencing (Fig. 4B-D, Extended Data Fig. 5).

Finally, we noted with interest that one of the cases found to harbor colibactin-producing bacteria was also the case identified with 8 distinct ERG+ PIA lesions (case 2, Fig. 3B). We therefore questioned whether bacterial genotoxins such as colibactin, along with inflammation, can contribute to genomic damage that promotes the formation of TMPRSS2:ERG gene fusions. We exposed LNCaP cells to the *E*. *coli* strain DH10B hosting a bacterial artificial chromosome (BAC) bearing the pks island that produces colibactin (PKS+), or hosting the empty pBeloBAC11 vector (PKS-) as previously described^13^ with or without the addition of TNFα to simulate inflammation (Fig. 4E) and then assessed dsDNA breaks by comet assay. Transient treatment of LNCaP cells with the PKS+ colibactin-producing bacteria with or without the addition of TNFα induced dsDNA breaks at or greater than that of 8 Gy of ionizing radiation at 4 hr and 24 hr after exposure (Fig. 4F). A similar trend was observed for induction of phosphorylated H2A histone family member X (γH2AX), as an indicator of dsDNA break repair (Fig. 4G).

In summary, we submit that our findings represent evidence in human tissues that bacterial infections drive early prostate cancer development. We speculate that the inflammation induced by infection, in combination with DNA damage induced by bacterial genotoxins, contributes to the development of precancerous lesions and oncogenic gene fusions, and promotes early prostate carcinogenesis (Extended Data Movie 1). We did not find evidence of persistent bacterial presence within prostate cancer however, as bacteria were never observed within cancerous regions in the cases in our 16S RISH and LPS and LTA IHC assays (Extended Data Fig. 4). This represents an important epidemiologic challenge in linking prostate infections to prostate cancer risk, as the initiating infection likely often occurs and is cleared many years prior to the cancer diagnosis.

Prostate infections and inflammation may contribute to other oncogenic events in addition to TMRPSS2:ERG gene fusions, and may contribute to carcinogenesis in ERG-cancers as well. It is of keen interest that we found ERG+ PIA and/or ERG+ LGPIN in apparent direct transition with ERG+ early invasive adenocarcinoma in the absence of any ERG+ HGPIN. Currently, the most accepted direct precursor to prostate cancer is HGPIN^29^, and it is hypothesized that PIA serves as a risk factor lesion that can directly transition to HGPIN^30^. It has also been previously hypothesized that prostate atrophy may give rise to carcinoma directly, and our current study would support this hypothesis in at least a subset of cases^31-33^. Whereas not all ERG+ precursor lesions may progress to invasive cancer, our finding of multiple foci of ERG+ precursors in infected cases would be in line with the multifocal nature of prostate cancer^34^. Overall, our study suggests that bacterial prostatitis should be considered as a legitimate risk factor for prostate carcinogenesis, and prompts the development of methodologies to detect undiagnosed prostate infections, as well as to mitigate infections and inflammation in the prostate, as a prostate cancer prevention strategy.

## Acknowledgments

We thank Jessica Hicks for assistance with IHC, Tracy Jones for assistance with slide scanning and block requests. We also thank Dr. Srinivasan Yegnasubramanian, Dr. William Nelson, Ajay Vaghasia, Dr. Cynthia Sears, and Dr. James Eshleman for helpful advice and discussion. We acknowledge Dr. Tim Phelps and Dr. David Rini for important input and guidance on Movie S1. We acknowledge and thank Dr. Jean*-*Philippe Nougayrède for providing the PKS+ and PKS-*E*. *coli* strains.

## Author contributions

K.S.S. conceived the study and all authors contributed to designing the experiments. K.S.S., E.S., J.C., L.M., S.E.E., Y.B., J.P.M. and C.J.H. performed the experiments and B.O. and A.M.D. performed the pathology assessments. A.K.M and M.C.H. assisted with developing the FISH assay. W.G. developed Movie S1. E.S. and K.S.S. wrote the manuscript and all authors assisted with editing the manuscript. All authors contributed to the analysis and interpretation of the data and approved the final manuscript.

## Data and materials availability

All data are available in the main text or the supplementary materials.

## References

1. Tomlins, S.A., et al. Recurrent fusion of TMPRSS2 and ETS transcription factor genes in prostate cancer. Science 310, 644–648 (2005).

2. Mosquera, J.-M., et al. Prevalence of TMPRSS2-ERG fusion prostate cancer among men undergoing prostate biopsy in the United States. Clinical cancer research : an official journal of the American Association for Cancer Research 15, 4706–4711 (2009).

3. Mani, R.S., et al. Inflammation-induced oxidative stress mediates gene fusion formation in prostate cancer. Cell Reports 17, 2620–2631 (2016).

4. Cerveira, N., et al. TMPRSS2-ERG Gene Fusion Causing ERG Overexpression Precedes Chromosome Copy Number Changes in Prostate Carcinomas, Paired HGPIN Lesions. Neoplasia 8, 826–832 (2006).

5. Klezovitch, O., et al. A causal role for ERG in neoplastic transformation of prostate epithelium. 105, 2105–2110 (2008).

6. Tomlins, S.A., et al. Role of the TMPRSS2-ERG gene fusion in prostate cancer. Neoplasia 10, 177–188 (2008).

7. Zong, Y., et al. ETS family transcription factors collaborate with alternative signaling pathways to induce carcinoma from adult murine prostate cells. Proceedings of the National Academy of Sciences of the United States of America 106, 12465–12470 (2009).

8. King, J.C., et al. Cooperativity of TMPRSS2-ERG with PI3-kinase pathway activation in prostate oncogenesis. Nature genetics 41, 524–526 (2009).

9. Carver, B.S., et al. Aberrant ERG expression cooperates with loss of PTEN to promote cancer progression in the prostate. Nature genetics 41, 619–624 (2009).

10. De Marzo, A.M., et al. Inflammation in prostate carcinogenesis. Nat Rev Cancer 7, 256-269 (2007).

11. Sfanos, K.S., Yegnasubramanian, S., Nelson, W.G. & De Marzo, A.M. The inflammatory microenvironment and microbiome in prostate cancer development. Nat Rev Urol 15, 11-24 (2018).

12. Nougayrède, J.-P., et al. Escherichia coli induces DNA double-strand breaks in eukaryotic cells. Science 313, 848–851 (2006).

13. Cuevas-Ramos, G., et al. Escherichia coli induces DNA damage in vivo and triggers genomic instability in mammalian cells. Proceedings of the National Academy of Sciences 107, 11537–11542 (2010).

14. Wilson, M.R., et al. The human gut bacterial genotoxin colibactin alkylates DNA. Science 363, eaar7785 (2019).

15. Xue, M., et al. Structure elucidation of colibactin and its DNA cross-links. Science 365, eaax2685 (2019).

16. Li, Z.-R., et al. Macrocyclic colibactin induces DNA double-strand breaks via copper-mediated oxidative cleavage. Nature Chemistry 11, 880–889 (2019).

17. Dziubanska-Kusibab, P.J., et al. Colibactin DNA-damage signature indicates mutational impact in colorectal cancer. Nature Medicine (2020).

18. Pleguezuelos-Manzano, C., et al. Mutational signature in colorectal cancer caused by genotoxic pks+ E. coli. Nature (2020).

19. Young, A., et al. Correlation of urine TMPRSS2:ERG and PCA3 to ERG+ and total prostate cancer burden. American Journal of Clinical Pathology 138, 685–696 (2012).

20. Morais, C.L., et al. ERG and PTEN status of isolated high-grade PIN occurring in cystoprostatectomy specimens without invasive prostatic adenocarcinoma. Human Pathology 55, 117–125 (2016).

21. Furusato, B., et al. ERG oncoprotein expression in prostate cancer: clonal progression of ERG-positive tumor cells and potential for ERG-based stratification. Prostate Cancer And Prostatic Diseases 13, 228 (2010).

22. Perner, S., et al. TMPRSS2-ERG fusion prostate cancer: an early molecular event associated with invasion. The American journal of surgical pathology 31, 882–888 (2007).

23. Haffner, M.C., et al. Molecular evidence that invasive adenocarcinoma can mimic prostatic intraepithelial neoplasia (PIN) and intraductal carcinoma through retrograde glandular colonization. J Pathol 238, 31–41 (2016).

24. McNeal, J.E. Prostatic microcarcinomas in relation to cancer origin and the evolution to clinical cancer. Cancer 71, 984–991 (1993).

25. van Leenders, G.J., et al. Intermediate cells in human prostate epithelium are enriched in proliferative inflammatory atrophy. Am J Pathol 162, 1529–1537 (2003).

26. Liu, X., et al. Low CD38 identifies progenitor-like inflammation-associated luminal cells that can initiate human prostate cancer and predict poor outcome. Cell Rep 17, 2596–2606 (2016).

27. Arthur, J.C., et al. Intestinal inflammation targets cancer-inducing activity of the microbiota. Science 338, 120–123 (2012).

28. Krieger, J.N., Dobrindt, U., Riley, D.E. & Oswald, E. Acute Escherichia coli prostatitis in previously health young men: bacterial virulence factors, antimicrobial resistance, and clinical outcomes. Urology 77, 1420–1425 (2011).

29. De Marzo, A.M., Haffner, M.C., Lotan, T.L., Yegnasubramanian, S. & Nelson, W.G. Premalignancy in prostate cancer: Rethinking what we know. Cancer prevention research (Philadelphia, Pa.) 9, 648–656 (2016).

30. Putzi, M.J. & De Marzo, A.M. Morphologic transitions between proliferative inflammatory atrophy and high-grade prostatic intraepithelial neoplasia. Urology 56, 828-832 (2000).

31. De Marzo, A.M., Marchi, V.L., Epstein, J.I. & Nelson, W.G. Proliferative inflammatory atrophy of the prostate: implications for prostatic carcinogenesis. Am J Pathol 155, 1985-1992 (1999).

32. Franks, L.M. Atrophy and hyperplasia in the prostate proper. The Journal of pathology and bacteriology 68, 617–621 (1954).

33. Liavåg, I. Atrophy and regeneration in the pathogenesis of prostatic carcinoma. Acta Pathologica Microbiologica Scandinavica 73, 338–350 (1968).

34. Andreoiu, M. & Cheng, L. Multifocal prostate cancer: biologic, prognostic, and therapeutic implications. Human Pathology 41, 781–793 (2010).

