## Supplemental Materials for "Oncogenic Gene Fusions in Non-Neoplastic Precursors as Evidence that Bacterial Infection Initiates Prostate Cancer"

**Materials and Methods**

Clinical Specimens

All specimens were obtained and studied under a Johns Hopkins Medicine Institutional Review Board (IRB) approved protocol. We searched the Johns Hopkins Hospital radical prostatectomy pathology reports from January 2016 through January 2018 for the terms prostatitis/cystitis, moderate or extensive acute inflammation, and granuloma. Hematoxylin and eosin (H&E) slides were reviewed to confirm that the degree/extent of inflammation contained in the radical prostatectomy specimens was highly atypical (see Extended Data Fig. 1). As these potential infections were undiagnosed, we cannot comment on when or how the infection occurred, and it is possible that the infection was from the diagnostic biopsy.

16S rRNA, *clbB*, and *ERG* RNA *in situ* Hybridization (RISH)

RISH was performed on formalin fixed paraffin embedded (FFPE) tissues using the Advanced Cell Diagnostics (ACD) RNAscope 2.5 HD Assay (ACD, Newark, CA, Cat. No. 322370). The probes used for the RISH assays were as follows: (1) 16S rRNA (Cat. No. 427731), (2) colibactin (*clbB*, Cat. No. 561691), (3) *ERG* (ACD, Cat. No. 604028), (4) peptidyl prolyl isomerase B (PPIB, positive control, Cat. No. 313901), (5) *Zea mays* superal1 (Sal1, negative control, Cat. No. 316381). RISH was performed according to a modified manufacturer's suggested protocol. Briefly, the slides were deparaffinized on a 60 °C heat block followed by xylene treatments. Slides were then washed in absolute alcohol and air dried followed by hydrogen peroxidase blocking treatment at RT for 10 minutes. The slides were treated with the antigen retrieval buffer for 15 minutes using the steaming method (temperature >= 99 °C) followed by a brief rinse with deionized water ($dH_2O$). Two enzyme treatment steps with lysozyme and achromopeptidase were performed exclusively in RISH targeting 16S rRNA to digest the cell wall of bacteria. Slides are first treated with lysozyme at 10 mg/ml (Sigma Aldrich, St. Louis, MO) followed by treatment with achromopeptidase at 30U/ml (Sigma Aldrich) for 10 minutes at 37 °C. Slides were then digested using protease plus pretreatment for 30 minutes at 40 °C. Finally, pre-warmed probes (40 °C) were hybridized onto the slides with 2 hours incubation at 40 °C.

The slides were washed using wash buffer and then the amplification steps were performed in the following order: amplification buffer 1, amplification buffer 2, amplification buffer 3, and amplification buffer 4 for alternating 30 minutes and 15 minutes at 40 °C with wash buffer between each amplification step. Then, amplification 5 was performed for 60 minutes at room temperature (RT) followed by a brief wash and amplification 6 for 30 minutes at RT. After a brief wash, a 1:1 solution of DAB A and B was applied to the tissue and incubated for 10 minutes at RT. This was then rinsed with $dH_2O$ and counterstained with hematoxylin for 2 minutes. The slides were rinsed, dehydrated, and coverslipped.

LPS and LTA immunohistochemistry (IHC)

IHC was performed manually with the following antibodies: (1) Lipotechoic Acid (LTA, Thermo Fisher Scientific, Waltham, MA, Cat. No. MA1-7402, Clone G43J) and (2) Lipopolysaccharide (LPS, Abcam, Cambridge, UK, Cat. No. 35654, Clone 2D7/1). Slides were deparaffinized at 60°C for 10 minutes and xylene treatment followed by rehydration in ethanol gradient and a rinse in $dH_2O$. After quick rinses in 0.1% TWEEN in $dH_2O$ solution and citrate buffer (Vector Laboratories, Burlingame, CA, Cat. No. H-3300), antigen retrieval was conducted using steaming method in EDTA buffer (Thermo Fisher Scientific, Cat. No. BP2473500) for 45 minutes for LPS antibody and HTTR buffer (Agilent Technologies, Santa Clara, CA, Cat. No. S169984-2) for LTA antibody. The slides were rinsed with TBST and blocking step was conducted using Dako REAL peroxidase blocking solution (Agilent Technologies, Cat. No. S202386-2) for 5 minutes at RT. The primary antibody was applied at 1:100 for LPS and 1:25 for LTA antibodies for 45 minutes at RT or overnight at 4°C, respectively. After a rinse, anti-mouse secondary antibodies were applied (Leica Biosystems, Wetzlar, Germany, Cat. No. PV6110) for 40 minutes at RT. The slides were rinsed and DAB (Sigma Aldrich, Cat. No. D4293) was applied for 20 minutes at RT. Counterstaining was done with Mayer's hematoxylin (Agilent Technologies, Cat. No. S330930-2) for 2 minutes and the slides were mounted with Cytoseal-60 (Thermo Fisher Scientific, Cat. No. 8310-16) after dehydration through an alcohol gradient.

FFPE DNA Extraction

DNA was extracted from macrodissected FFPE tissues from areas that were positive for both 16S rRNA RISH and LPS IHC in cases 2 and 12. DNA extraction was conducted using a phenol:chloroform-based method. Briefly, 5 µm tissue sections from areas with positive bacterial signature on adjacent cuts were macro-dissected using a sterile scalpel and placed in a 2 ml Eppendorf tube. The tissue was autoclaved to 120 °C for 25 minutes in an alkali digestion buffer (0.1 M NaOH in 1% SDS solution) for reversal of formalin-induced crosslinking. After a brief cooling of the tissue, 500 µl of 25:24:1 phenol:chloroform:isoamyl alcohol mixture was added. The mixture was agitated for 5 minutes at RT and centrifuged at 10,000 X g for 5 minutes at RT. The upper aqueous layer was transferred to a new tube. The agitation and centrifugation was repeated twice. Then, 1 volume of isopropanol and 0.1 volume of 3M sodium acetate was added to the aqueous layer and mixed. The mixture was then centrifuged at 10,000 X g for 30 minutes at RT. The supernatant was discarded, and the pellet was rinsed gently with 1 mL of 85% ethanol. The resultant washed pellet was air dried and re-suspended in molecular grade $dH_2O$.

Colibactin *clbB* Quantitative PCR (qPCR)

qPCR was conducted using a primer set (*32*) and probe targeting the *clbB* gene in the colibactin PKS island as follows: pks-F 5'- GCGCATCCTCAAGAGTAAATA-3', pks-R 5'-GCGCTCTATGCTCATCAACC-3', pks probe 5'-FAM-TATTCGACACAGAACAACGCCGGT-BHQ1-3' (probe designed by Dr. Julia Drewes in the laboratory of Dr. Cynthia Sears at Johns Hopkins). qPCR was performed using Bio-Rad iTaq Universal Probe Supermix (Bio-Rad, Hercules, CA) system following the manufacturer suggested protocol with 7-20 ng of input DNA. The cycling conditions for the assay were 95°C for 10 minutes followed by 40 cycles of 95°C for 15 seconds, 60°C for 1 minute, and a final extension at 72°C. qPCR was conducted on the Bio-Rad CFX Connect Real Timer PCR Detection System and the data was analyzed using Bio-Rad CFX Manager 3.1 software.
The amplified product from the qPCR was visualized via gel electrophoresis, gel extracted, and purified using the QiaQuick Gel Extraction kit (Qiagen, Hilden, Germany). The eluted products were then Sanger sequenced at the Johns Hopkins Genetic Resources Core Facility and aligned with bacterial genomes using NCBI BLAST.

16S rDNA PCR

DNA extracted from Case 2 and Case 12 was amplified via PCR with the following primer sets designed as universal primers against the 16S rRNA gene: 27F 5'-AGAGTTTGATCMTGGCTCAG-3' + 519R 5'- GWATTACCGCGGCKGCTG-3', 533F 5'-GTGCCAGCAGCCGCGGTAA-3' + 907R 5'- CCGTCAATTCMTTTRAGTTT-3', and V6-F 5'- CAACGCGWRGAACCTTACC-3' and V6-R 5'-CRRCACGAGCTGACGAC-3'. The amplified product from the PCR was visualized via gel electrophoresis and purified using the QiaQuick PCR Purification kit (Qiagen). The eluted products were then Sanger sequenced at the Johns Hopkins Genetic Resources Core Facility and aligned with bacterial genomes using NCBI BLAST.

ERG and PIN4 IHC

IHC was performed on the Ventana Discovery Ultra IHC/ISH system (Roche Diagnostics, Basel, Switzerland) with the following antibodies: (1) ERG (Roche Diagnostics, Cat. No. 790-4576, Clone EPR3864) and (2) PIN4 – Cytokeratin1/5/10/14 (Enzo Life Sciences, Farmingdale, NY, Cat. No. ENZ-C34903), p63 (Biocare Medical, Pacheco, CA, Cat. No. SKU: 163, Clone 4A4) and AMACR (Zeta Corporation, Arcadia, CA, Cat. No. Z2001, Clone 13H4). IHC was performed per the manufacturer's protocol. The slides were steamed for 32 minutes (for ERG) and 48 minutes (for PIN4) in Cell Conditioning 1 (CC1) solution (Roche Diagnostics, Cat. No. 950-124) for antigen retrieval. Then the corresponding primary antibodies were applied with the following conditions: ERG (pre-diluted) for 32 minutes at RT and PIN 4 – a combination of 1:50 dilution of Cytokeratin 1/5/10/14 and P63 for 40 minutes at RT followed by 1:50 dilution of AMACR for 32 minutes at RT. The Discover HQ HRP hapten-linked multimer detection kit (Roche Diagnostics, Cat. No. 760-4602) and the Discovery Amp HQ kit (Roche Diagnostics, Cat. No. 760-052) were used to develop ERG staining while the PIN4 staining was developed in the Discover HQ HRP hapten-linked multimer detection kit.

As a confirmation of the ERG staining, we also performed ERG IHC with a separate antibody (Biocare Medical, Concord, CA, SKU: 421, Clone 9FY) and manual staining with the Power Vision+ Poly-HRP IHC kit (Leica Biosystems, Cat. No PV6109).

TMPRSS2:ERG Fluorescent *in situ* Hybridization (FISH) and Imaging

FISH was conducted on FFPE radical prostatectomy tissues using the TMPRSS2-ERG (21q22) Deletion, Break Triple color FISH assay (Leica Biosystems, Cat. No. KI-10726). The assay was conducted using a modified manufacturer's method. FFPE tissues were deparaffinized at 60 °C for 10 minutes followed by 2X xylene treatment. The tissue was re-hydrated through an alcohol gradient and $dH_2O$. The slides were then treated with 0.2N HCl for 15 minutes at RT followed by antigen retrieval using a 10 mM sodium citrate solution (Vector Laboratories, Cat. No. H-3300) at 80 °C for 40 minutes. The slides were then treated with 2X SSC, $dH_2O$, and 0.2N HCl at RT for 2 minutes each. Protease digestion was conducted using ISH protease 3 (Roche Diagnostics, Cat. No. 780-4149) on the Benchmark Ultra IHC/ISH system (Roche Diagnostics) for 40 minutes at 37 °C. The slides were washed 2X in $dH_2O$ and fixed in 10% neutral buffered formalin (NBF). After 2X brief $dH_2O$ wash, the slides were dehydrated and air dried. The slides were stored at -20 °C until the probe hybridization.

The slides were warmed to 45 °C and 5 µl of the Leica TE break apart probe was added to the area of interest. A coverslip was placed over the probe and sealed. The probes were hybridized in a Thermobrite slide denaturation and hybridization system (Leica Microsystems, Cat. No. 23-021-580) at 80 °C for 5 minutes followed by 37 °C for 16-24 hours. Post hybridization, the coverslip was removed, and the slides were washed with pre warmed 2XSSC/0.3%IgePal CA-60 wash (Thermo Fisher Scientific, Cat. No. 15557044 and Sigma Aldrich, Cat. No. I8896, respectively) for 2 minutes at 72 °C. This was followed by a second wash of 2X SSC/0.1% IgePal CA-60 for 1 minute at RT. After a brief rinse in $dH_2O$, the slides were dehydrated and air dried. The tissue was counterstained with DAPI (Leica Biosystems, Cat. No. LK-095A) at a dilution of 1:5 in counterstain diluent (Leica Biosystems, LK-097A) and coverslipped. The slides were stored in 4 °C in the dark until they were imaged.

The slides were initially visualized using a Nikon Eclipse 2000 fluorescence microscope under 40X objective magnification with channels for FITC (green), Cy3 (red), and CFP (aqua) markers respectively. For individual cell analysis, the slides were imaged using the TissueFAXS Plus (Tissue Gnostics) automated microscopy workstation equipped with a Zeiss Z2 Axioimager microscope. First, each slide was imaged with a 10X objective with the DAPI channel to generate a preview image of the slide. Following the preview, specific regions of interest were identified and imaged using the 63X oil objective. The ranges of exposure time for each filter was determined for each individual sample: DAPI (50ms, 0-2500), FITC (250ms, 0-4095), Cy3 (175ms, 0-4095), ET-A (175ms, 0-4095). TissueFAXS Viewer (TissueGnostics, Wien, Austria) software was used to visualize the composite images and Photoshop (Adobe, San Jose, CA) was used to further reduce background levels. Assignment of TMPRSS2:ERG gene status in individual cells in areas of interest was conducted by two reviewers. Each region was assessed for a minimum of 100 non-overlapping distinct cells. Each cell was scored if both sets of telomeric TMPRSS2 (red) and centromeric ERG (blue) signals were present. If either of those signals were missing or a clear score was unable to be assigned, an unquantifiable score (*) was assigned to the cell. A region was defined as fusion positive if > 10% of the cells showed the same fusion pattern.

*In vitro* Cell Culture Modeling

LNCaP cells (obtained from ATCC, Manassas, VA, Cat. No. CRL-1740) were cultured in RPMI with 10% FBS in 6 well plates or T25 flask to 80% confluence. PKS producing bacteria were obtained courtesy of Dr. Jean-Phillipe Nougayrède. Briefly, for the *in vitro* assay *E. coli* strain DH10B hosting a bacterial artificial chromosome (BAC) bearing the pks island (PKS+) was used as colibactin producing strain while DH10B hosting the empty pBeloBAC11 vector (PKS-) was used as the negative bacterial control. PKS+ and PKS- bacteria were grown shaking overnight at 37°C in LB broth. Prior to the start of the experiment, the number of LNCaP cells per well/flask was measured in a proxy flask using an automated cell counter (Invitrogen Countess II). Fresh media was changed and PKS+/PKS- bacteria (MOI 100:1) and TNFα (100 ng/ml) was added to the cells. The treatment groups were no treatment control (NTC), TNFα only, PKS+ only, PKS- only, TNFα & PKS+, and TNFα & PKS-. At four hours post infection, cells were harvested for a 4H time point or the cells were washed with DPBS and media with gentamycin (20 µg/ml) was added to the cells. TNFα (100ng/ml) was added to the applicable groups. At 24 hours, the cells were washed and re-suspended in DPBS and used for COMET assay directly and stored at -20°C for western blot. Etoposide (10 µM) and ionizing radiation (8 Gy) treated cells were used as positive controls for DNA damage.

Comet Assay

Neutral comet assay was performed using Trevigen CometAssay kit (Trevigen, Gaithersburg, MD, Cat. No. 4250-050-K). Cells were washed with 1X DPBS, trypsinized for 5 minutes at 37 °C, washed, and re-suspended in 1X DPBS to a final concentration of $1x10^5$ cells per ml. The COMET assay was performed according to the manufacturer's protocol. Briefly, 50 µl of washed $1x10^5$ cells/ml were mixed with 500 µl LMA agarose at 37 °C. Next, 30 µl of the cell agarose mixture was put onto warmed Comet Slides to form a uniform layer which was set on a cool flat metal surface at 4 °C. Cell membrane lysis was conducted using the pre-cooled Trevigen lysis solution overnight at 4 °C. The slides were washed in 1X TBE buffer followed by gel electrophoresis at a fixed voltage of 21 V for 45 minutes at 4 °C. Slides were rinsed in $dH_2O$ and 70% ethanol and dried in a 37 °C incubator until the agarose dehydrated forming a flat surface (~2 hours). Then, the slides were stained with 1X SYBR green solution (Invitrogen, Cat. no. S7585) for 30 minutes at RT in the dark and visualized under a Nikon fluorescence microscope in the FITC channel. Images were taken using Roper scientific image software at 4X for analysis.

CometScore 2.0 was used to analyze the images from the Comet assay and the tail moment was used as a measure of DNA damage. The analyzer was blinded to the treatment groups. The software was used according to the developer's recommendations. A minimum of 50 individual comets were analyzed for each group. Statistical analysis was conducted individually for each biological replicate. A non-parametric one way ANOVA followed by post hoc analysis (Dunn's test) was used to compare the means of the different groups.

Western Blot

Cells from the *in vitro* assay were used for western blot. ɣH2AX protein was detected from whole protein isolate. Briefly, cells were harvested and lysed in using lysing buffer supplemented with phosphatase, benzoase, and proteinase inhibitors as per the manufacturer's protocol. The protein concentration was measured using a Pierce BCA assay (Thermo Fisher Scientific, Cat. No. 23225) and run on an 8% Bis-Tris SDS-PAGE gel (Thermo Fisher Scientific, Cat. No. NW00105BOX). The gel was transferred onto a nitrocellulose membrane and blocked with a BSA blocking buffer (LI-COR, Lincoln, NE, Cat. No. 927-50000). The membrane was stained with 1:1000 dilution of ɣH2AX (MilliporeSigma, Burlington, MA, Cat. No. 05-636, Clone JBW301) and 1:5000 dilution of β-Actin (Cell Signaling Technologies, Danvers, MA, Cat. No. 3700S, Clone 8H10D10) primary antibodies shaking overnight at 4°C. The membranes were washed and incubated in the anti-mouse secondary (LI-COR, Cat. No. 925-68070) shaking for 45 minutes at RT. The membranes were washed and visualized using an Odyssey scanner at 562 nm absorbance. Densitometry quantification was conducted using the ImageJ Gel analysis algorithm.

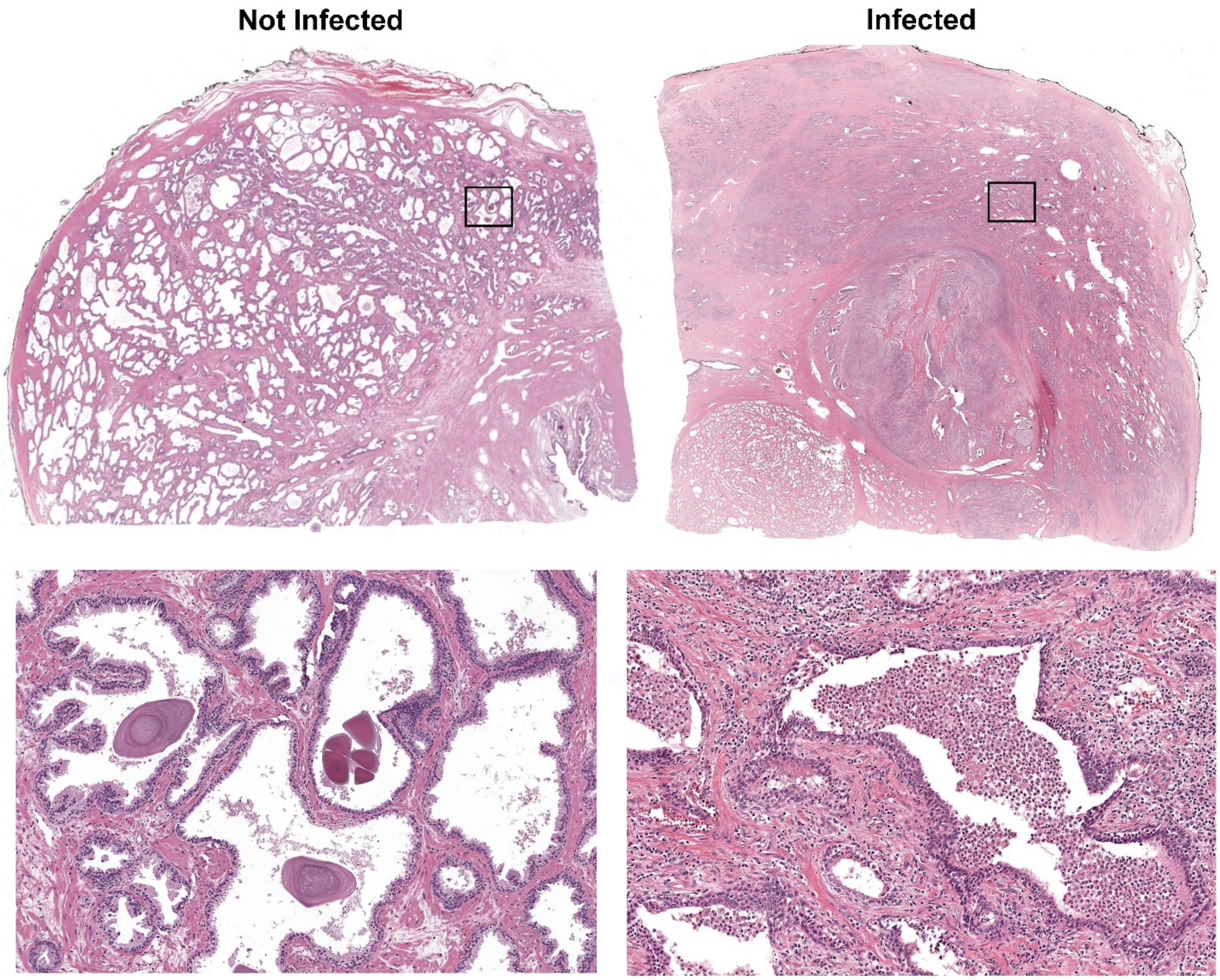


**Extended Data Fig. 1**. Example of a whole tissue section (top row) from a typical radical prostatectomy specimen (“non-infected”) versus an atypical, highly inflamed case (“infected”) suspicious for an active infection. The “infected” section stained positive for bacteria with 16S RISH, as well as by LPS IHC. Bottom images show magnified view of the boxed areas in the whole tissue sections (100X magnification).

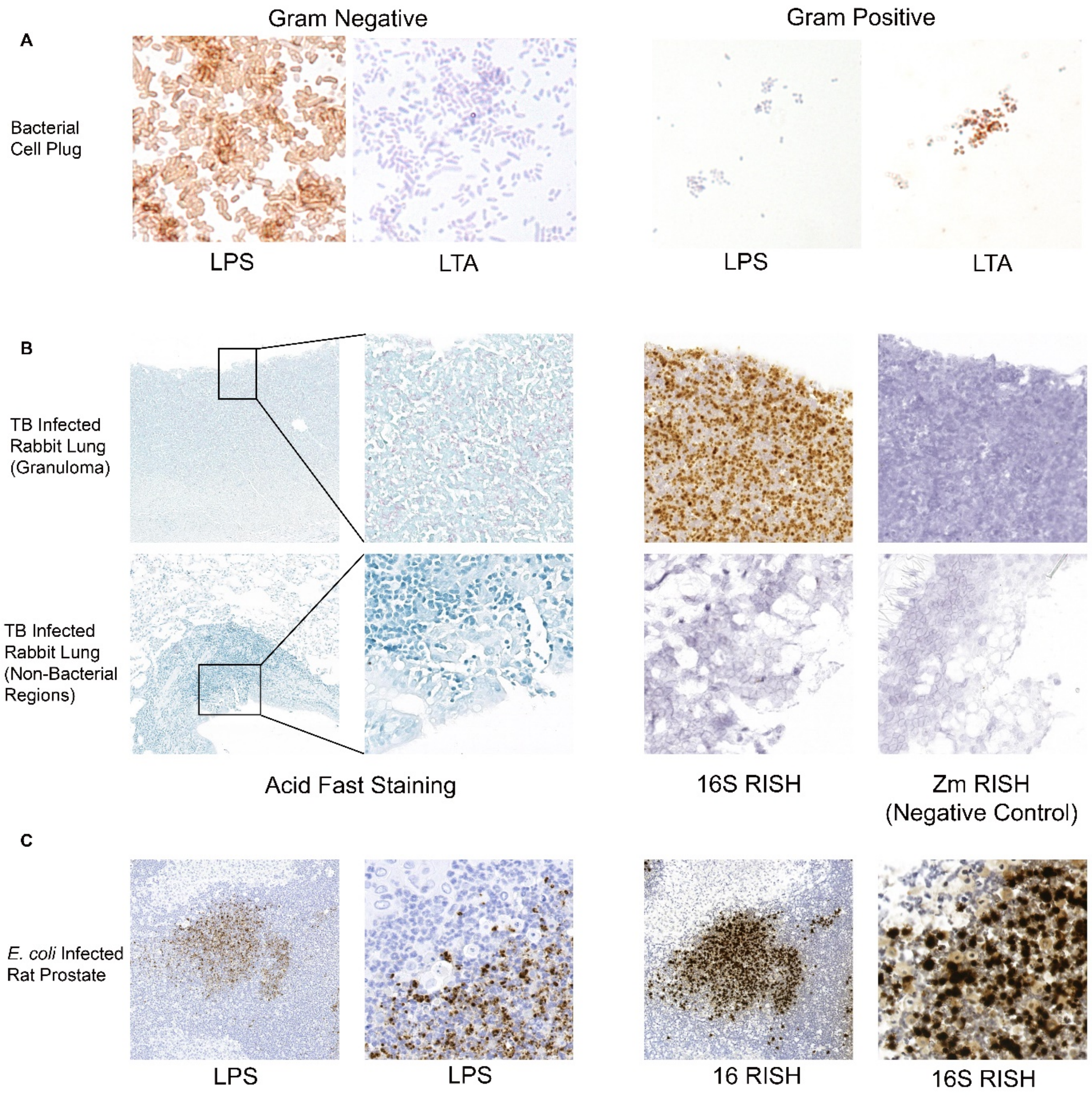


**Extended Data Fig. 2**. LPS and LTA IHC and 16S RISH assay validation. (A) Cell plugs of gram negative (*Klebsiella pneumoniae*) and gram positive (*Streptococcus* sp.) bacteria stained with the LPS and LTA IHC assays. (B) *Mycobacterium* (TB) infected rabbit lung stained with acid fast staining or 16S RISH. Only the acid fast positive areas are positive by 16S RISH. *Zea mays* (Zm) superal1 RISH serves as a negative control for the RISH assay. (C) *E. coli* infected rat prostate stained with LPS IHC and 16S RISH.

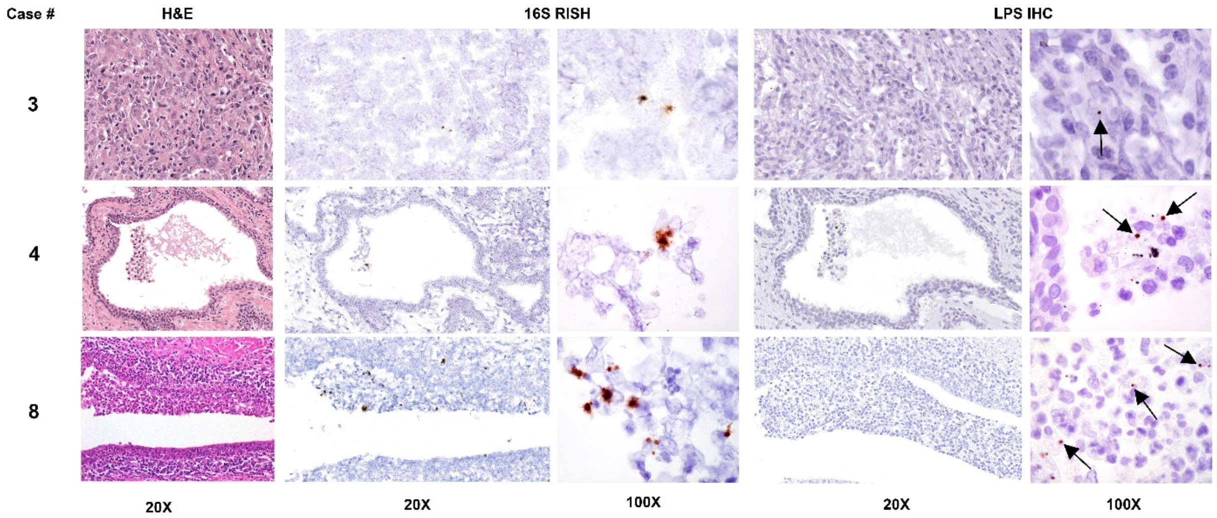


**Extended Data Fig. 3**. Additional examples of detection of bacteria in radical prostatectomy specimens from case 3, 4, and 8 with 16S RISH and LPS IHC. Objective magnification denoted. Arrows point to bacteria in association with immune cells. Images from case 3 show a region of severe inflammation obscuring epithelium. Images from case 4 and case 8 show PIA lesions.

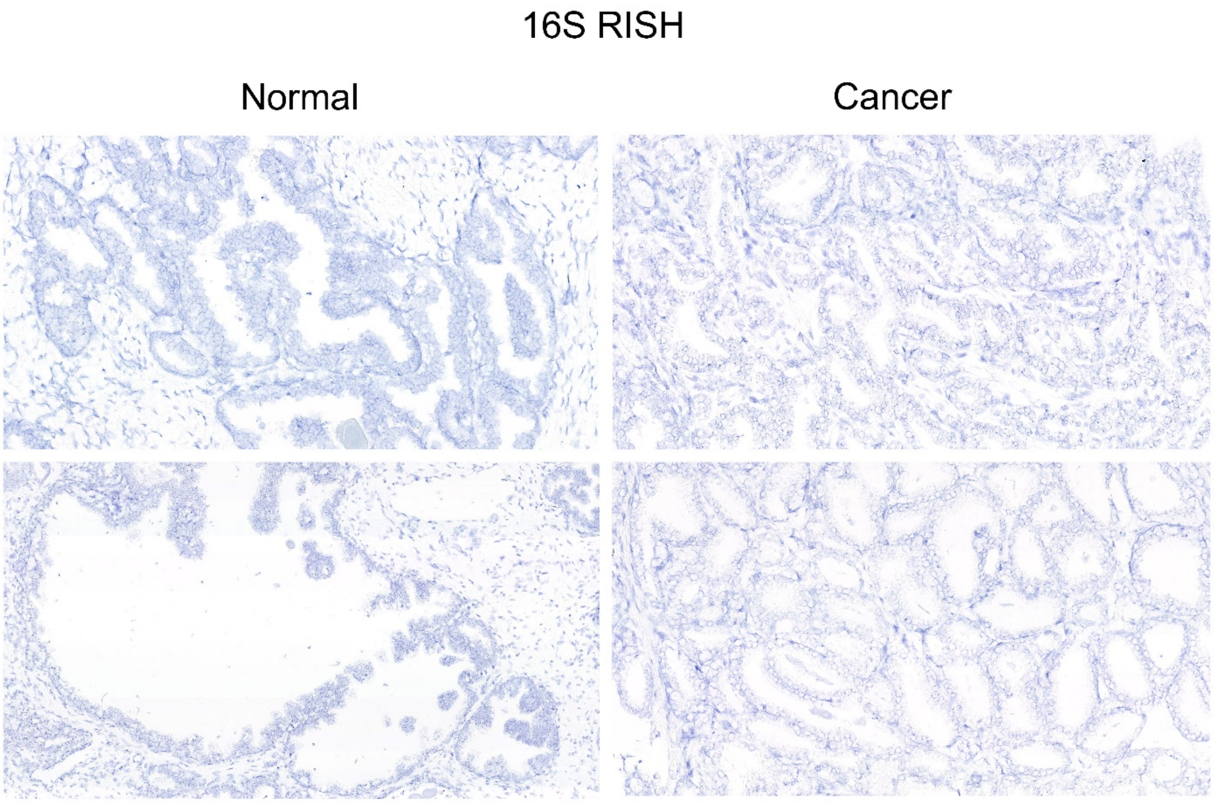


**Extended Data Fig. 4**. Bacterial signals were not observed by 16S RISH in normal-appearing (non-atrophic, not PIN), non-inflamed regions of prostate or in prostate cancer.

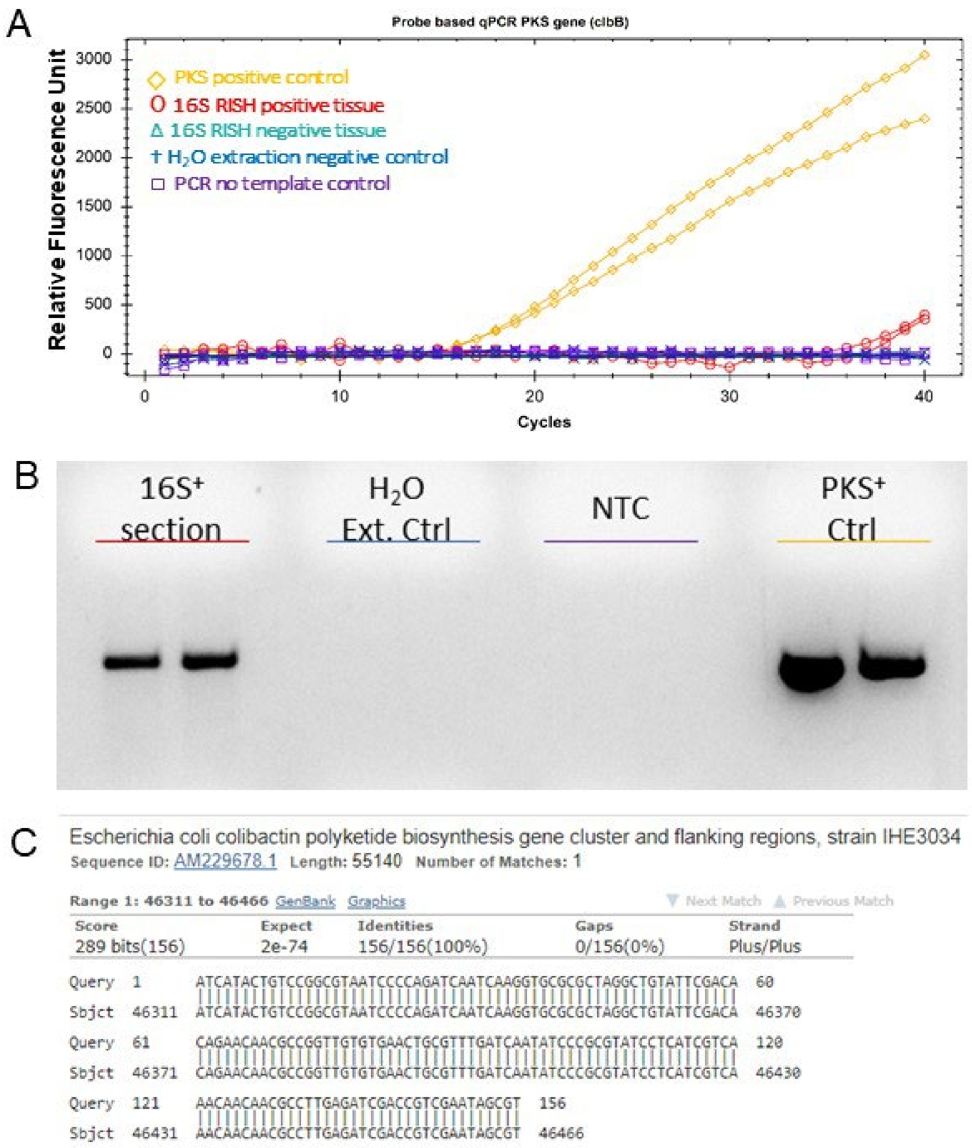


**Extended Data Fig. 5**. Detection of colibactin *clbB* gene in DNA extracted from case 12. (A) qPCR for *clbB*. (B) Agarose gel of qPCR products. (C) Alignment of sequenced qPCR product to the pks island encoding colibactin from *E. coli*.

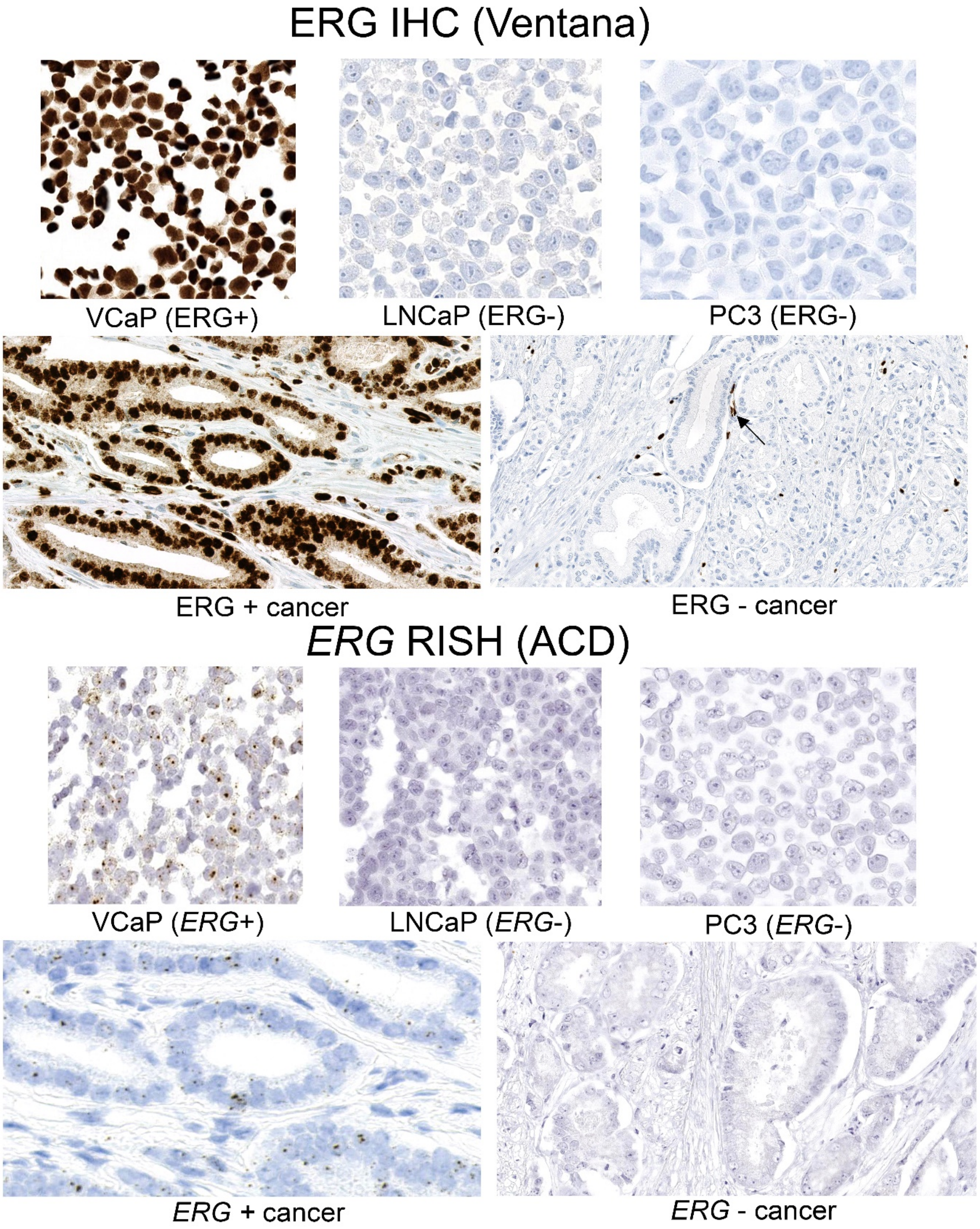


**Extended Data Fig. 6**. Validation of ERG IHC and *ERG* RISH assays using VCaP cells and ERG+ cancer as positive controls and LNCaP, PC3, and ERG- cancer as negative controls. Arrow points to ERG+ endothelial cells in a blood vessel as an internal positive control in an ERG- cancer.

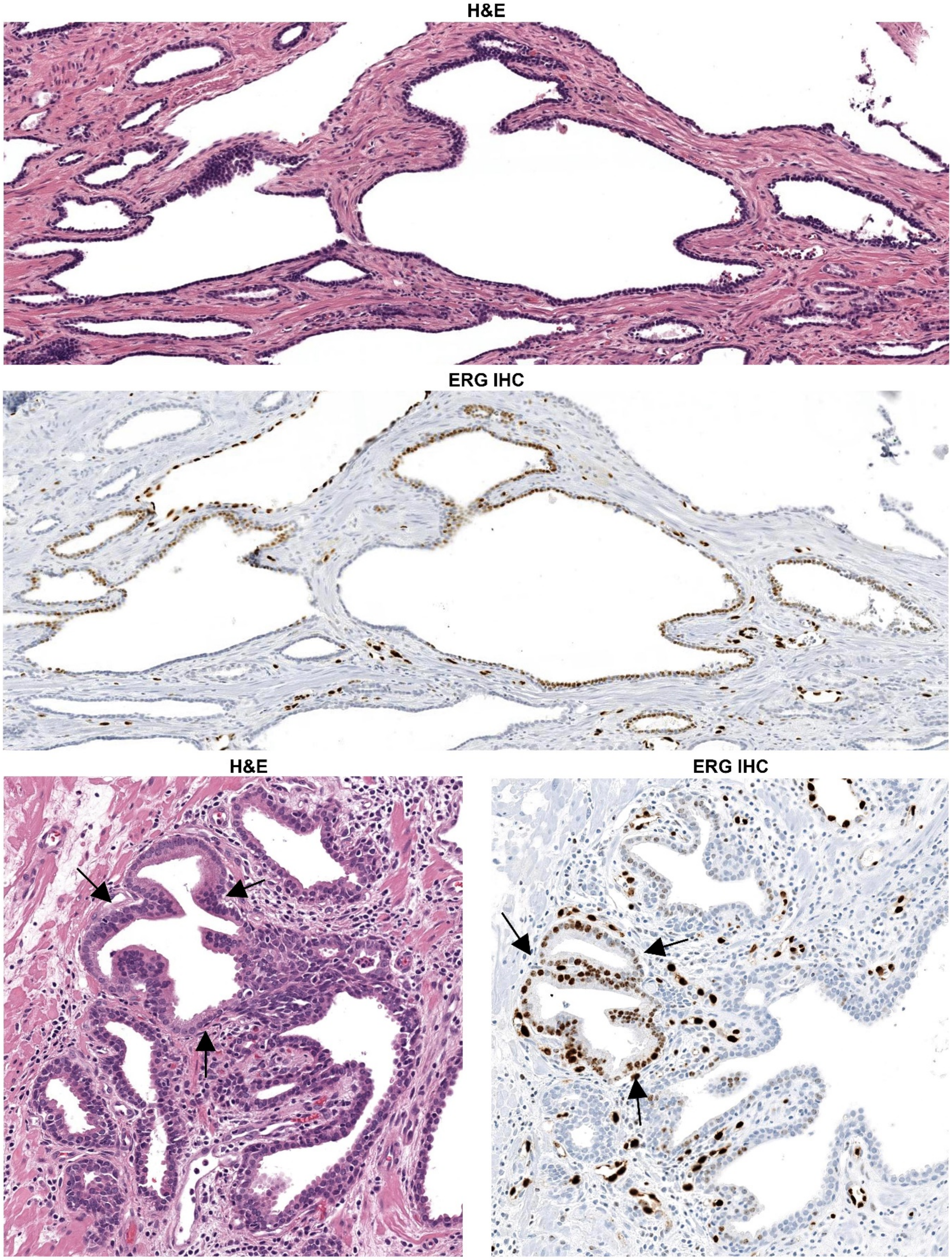


**Extended Data Fig. 7**. Examples of ERG+ PIA (top) and ERG+ PIA merging with LGPIN (bottom) identified in unselected cases. Arrows denote the LGPIN lesion.

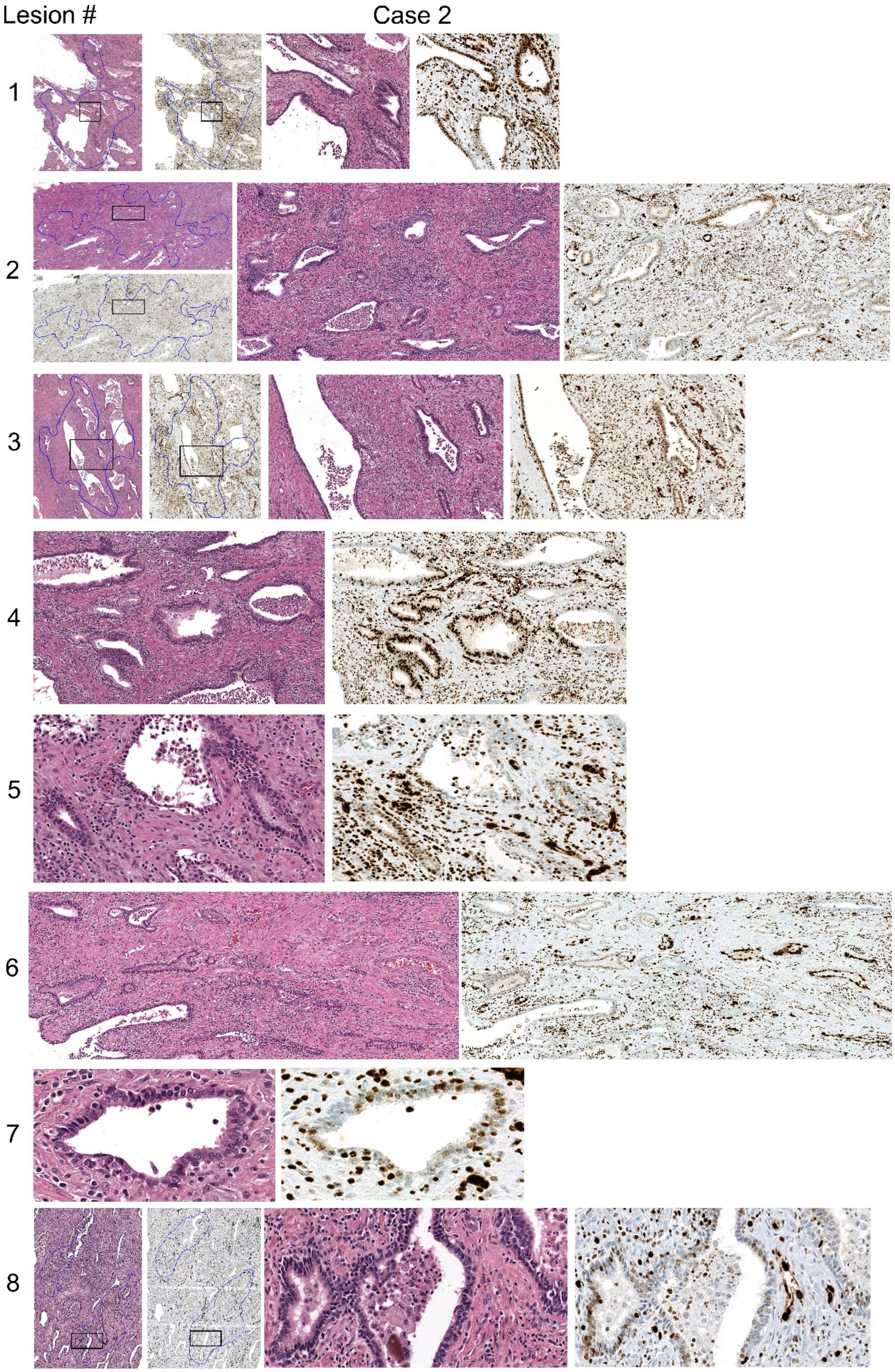
Lesion #
Case 2
1
2
3
4
5
6
7
8

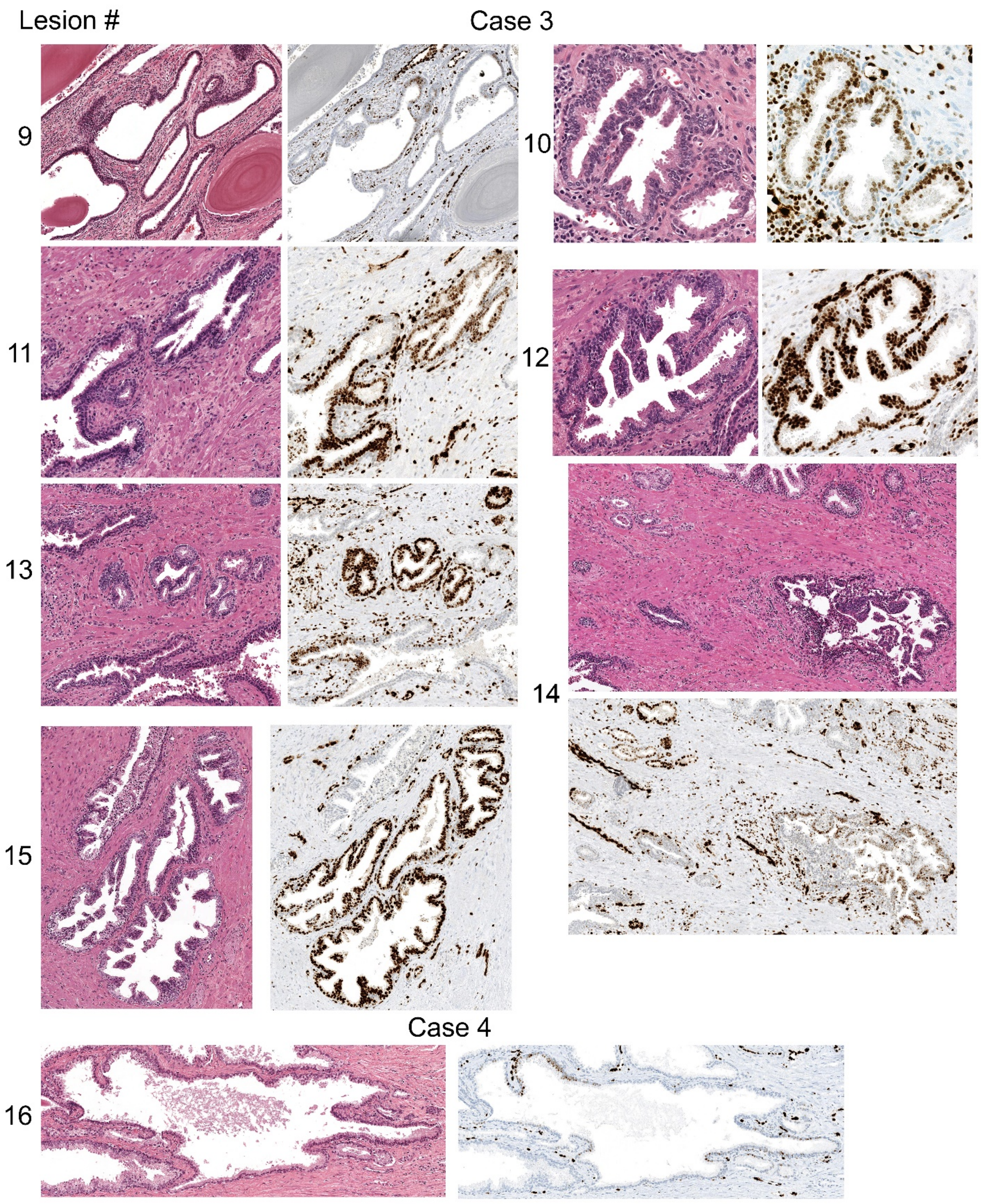
Lesion #
Case 3
9
10
11
12
13
14
15
Case 4
16

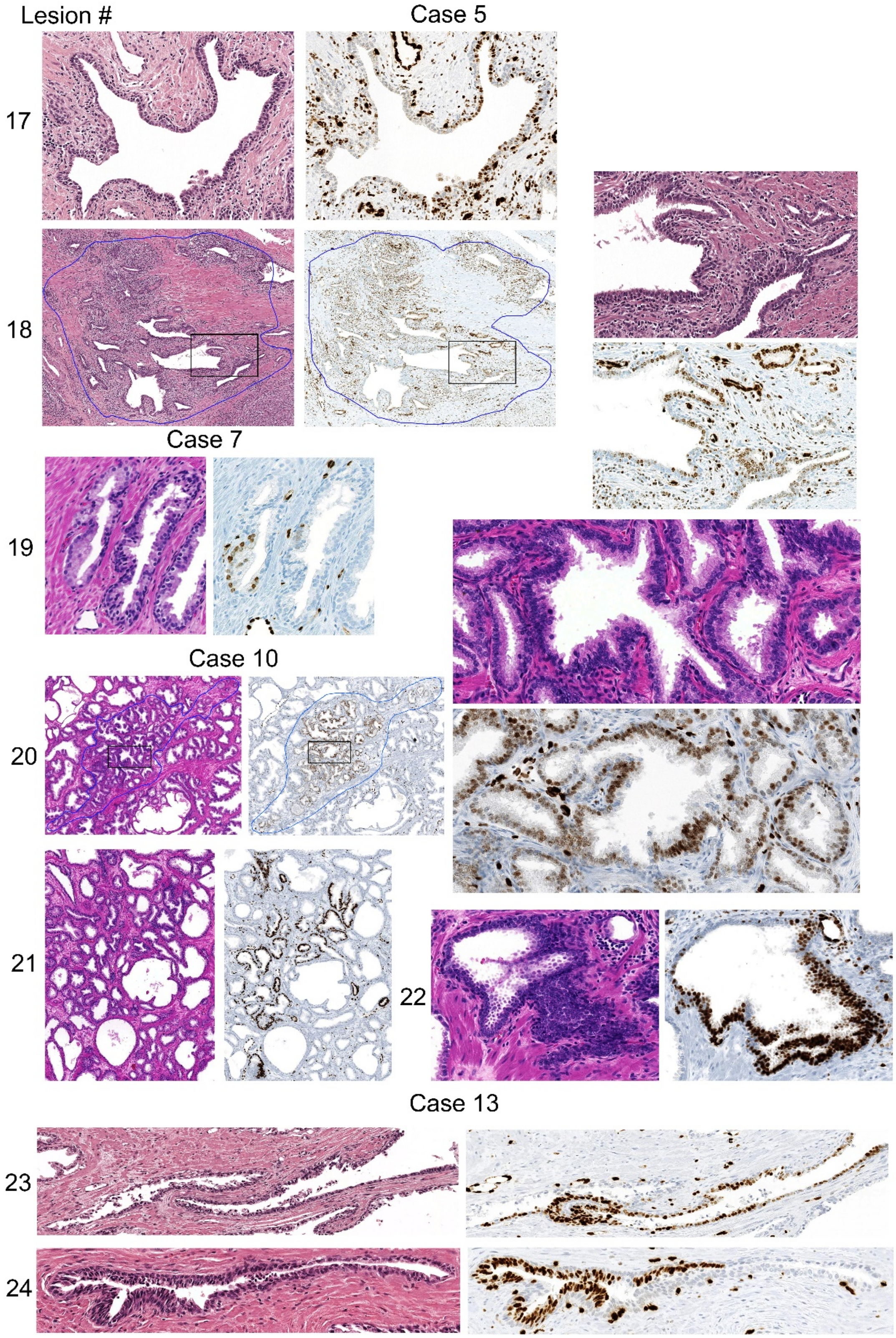


**Extended Data Fig. 8**. H&E and ERG IHC for all ERG+ PIA, LGPIN, and HGPIN foci identified in the study (lesion 1-24, see Table S2).

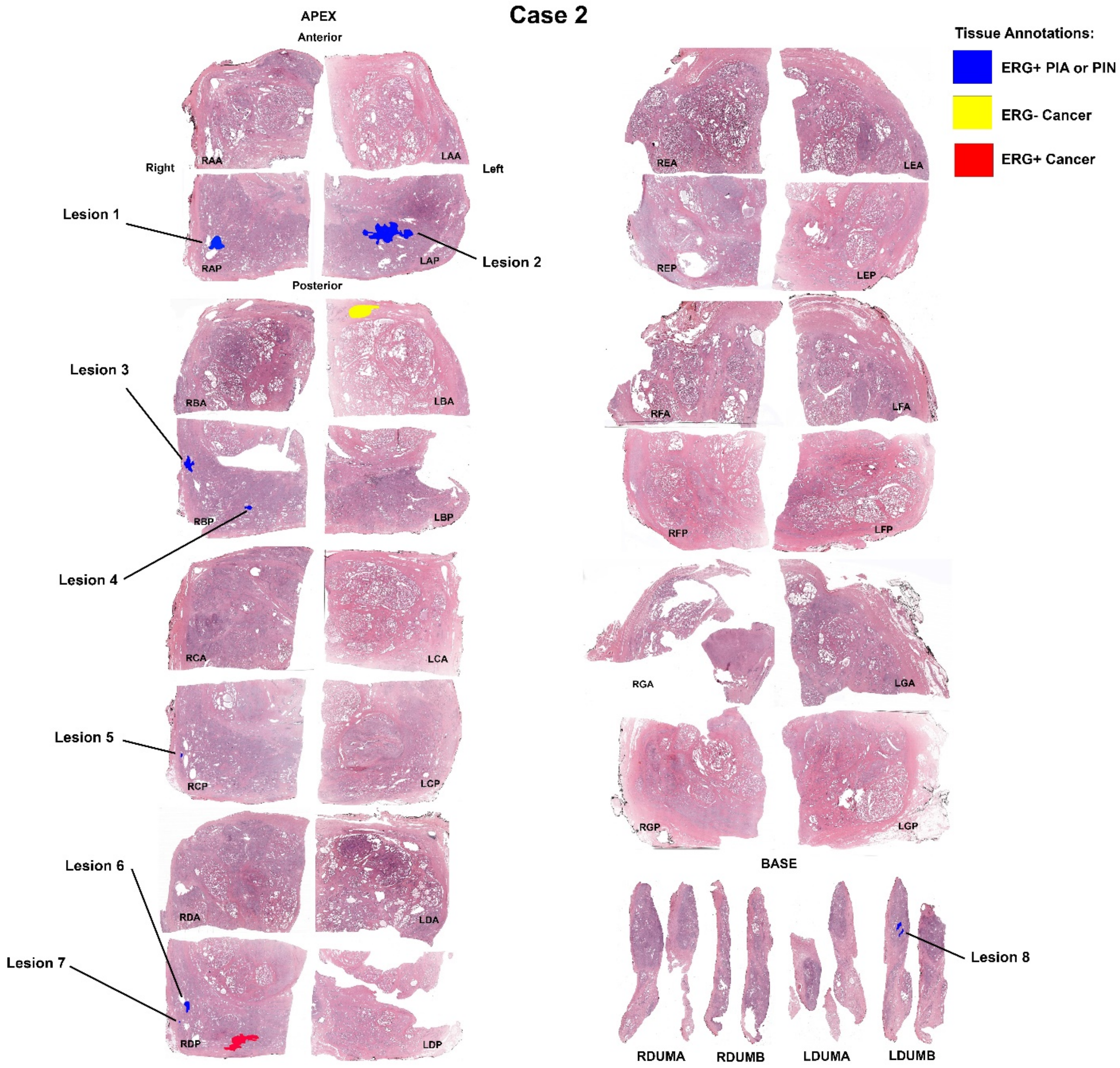


**Extended Data Fig. 9**. Prostate map showing location of ERG+ PIA (lesion 1-8) foci (blue), ERG- cancer (yellow), and ERG+ cancer (red) in case 2.

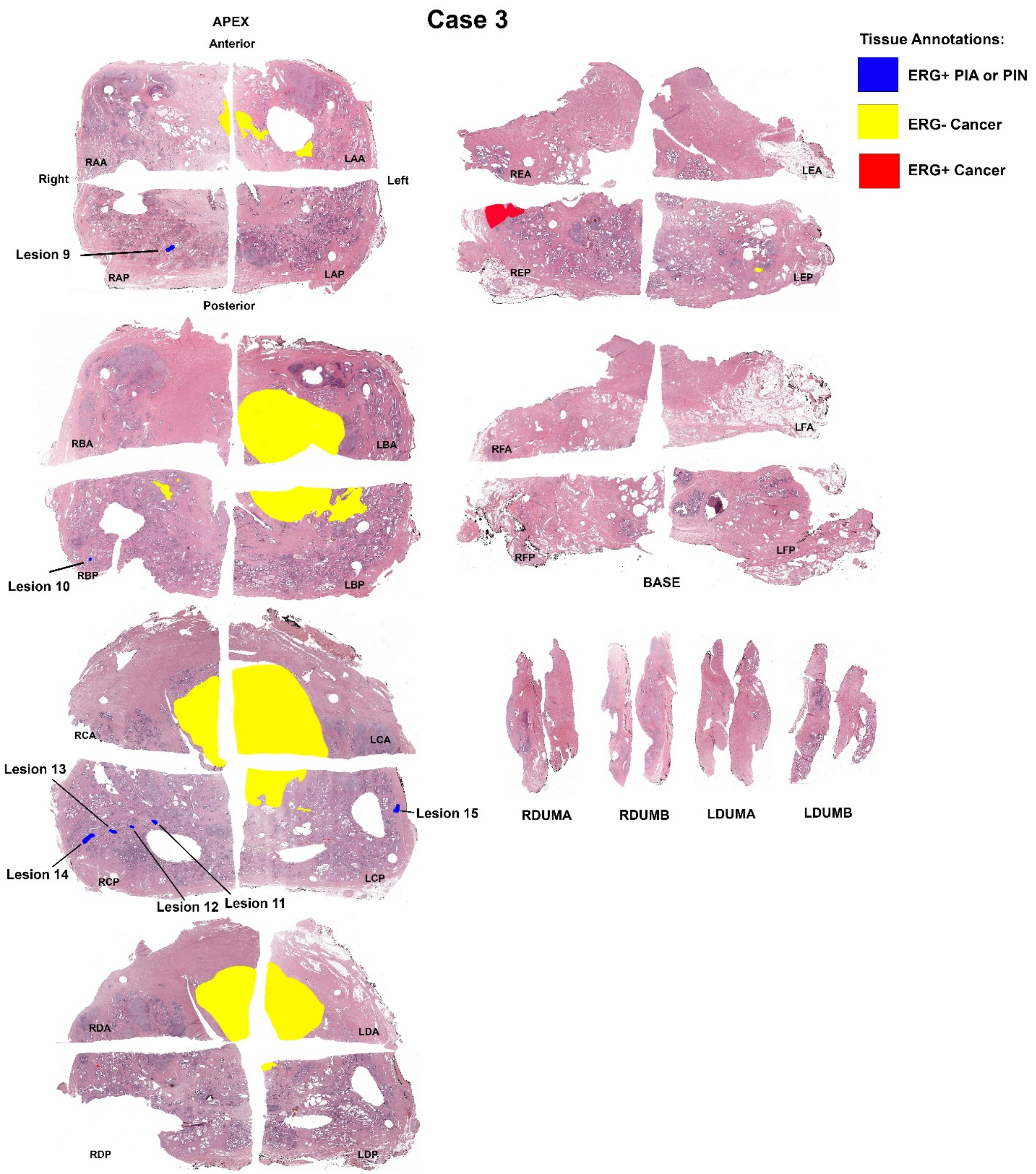


**Extended Data Fig. 10**. Prostate map showing location of ERG+ PIA (lesion 9), LGPIN (lesion 10-14), or HGPIN (lesion 15) foci (blue), ERG- cancer (yellow), and ERG+ cancer (red) in case 3.

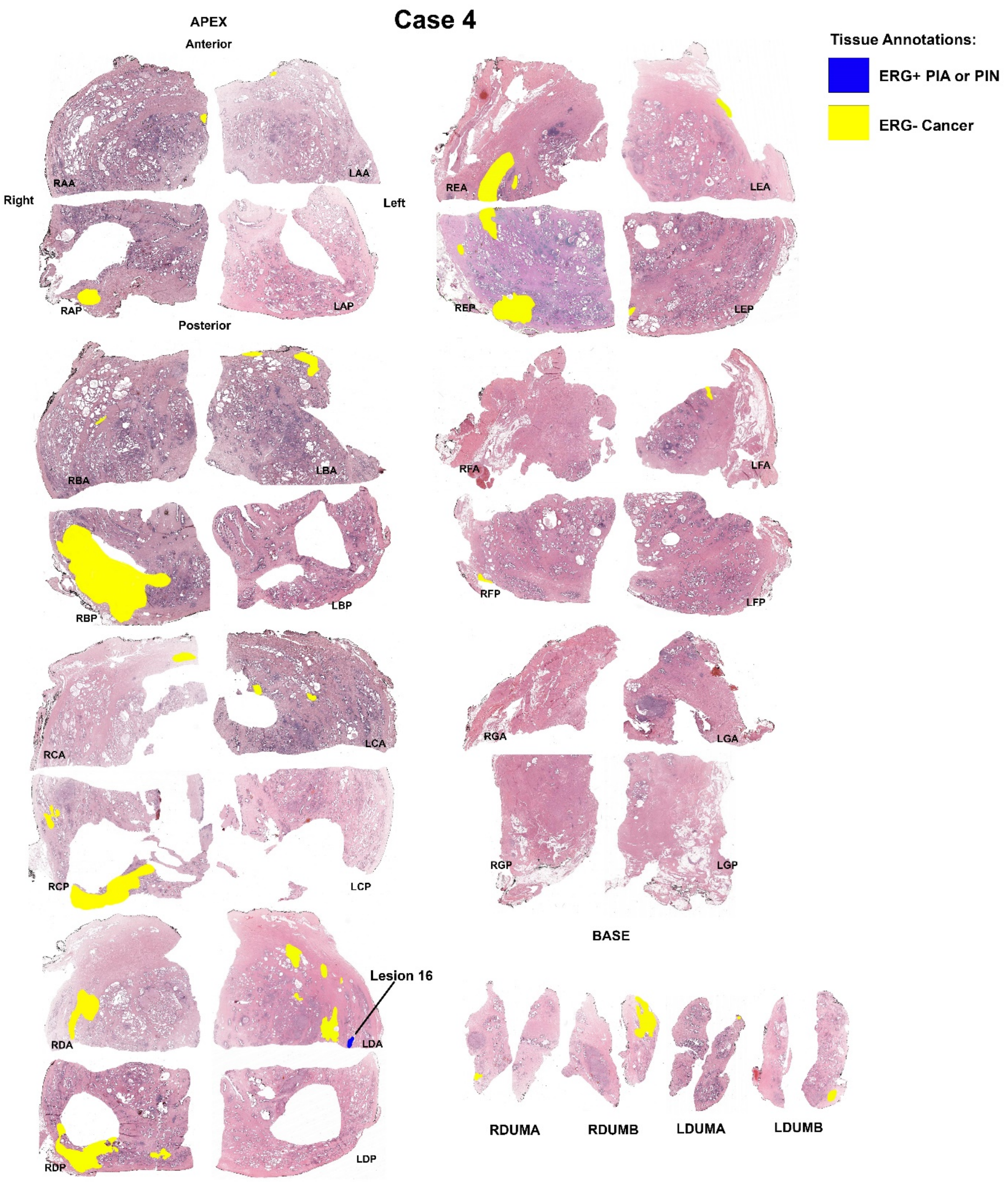


**Extended Data Fig. 11**. Prostate map showing location of ERG+ PIA merging with LGPIN/HGPIN (lesion 16) foci (blue) and ERG- cancer (yellow) in case 4. There was no ERG+ cancer found in this case.

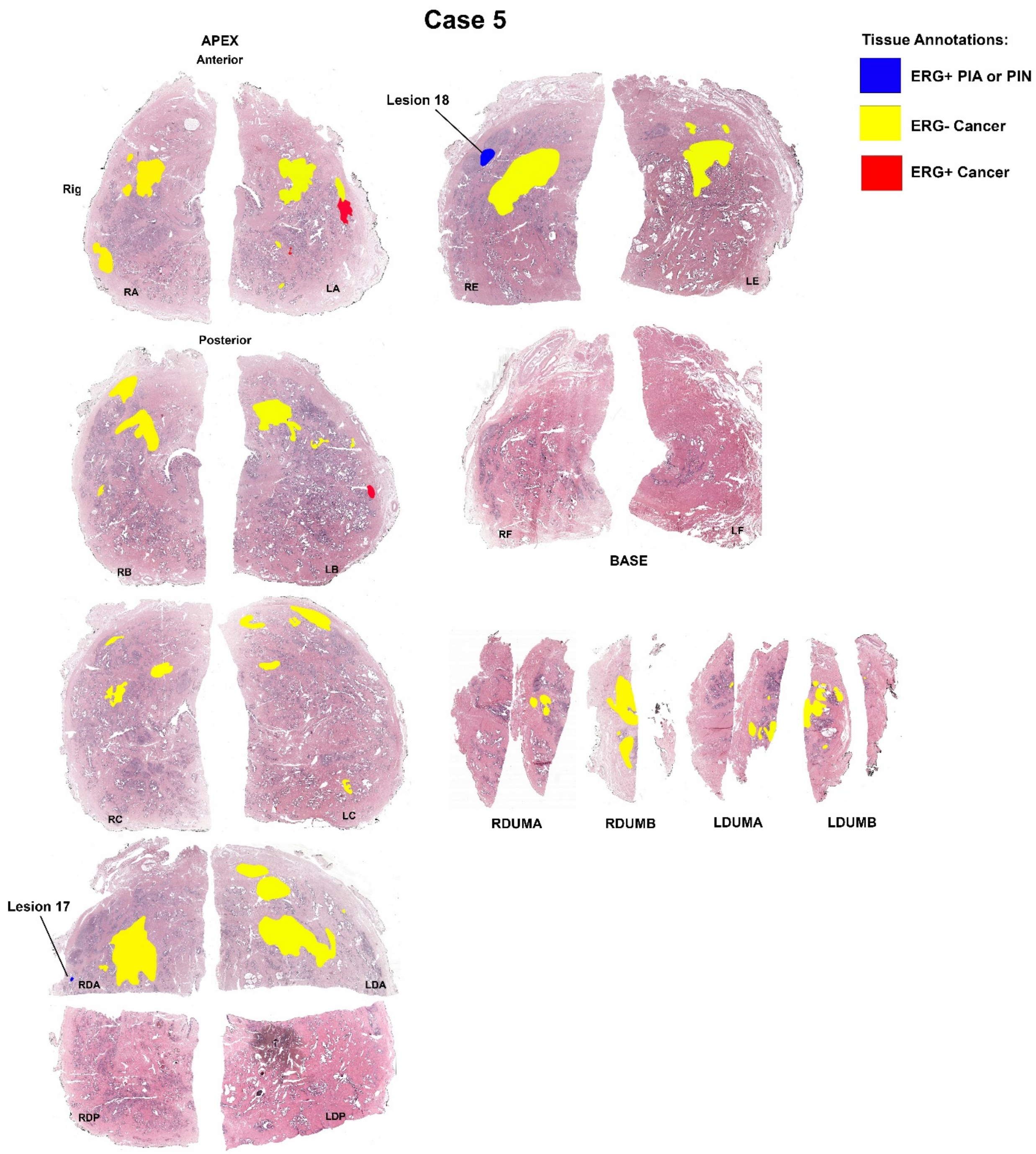


**Extended Data Fig. 12**. Prostate map showing location of ERG+ PIA (lesion 17 and 18) foci (blue), ERG- cancer (yellow), and ERG+ cancer (red) in case 5.

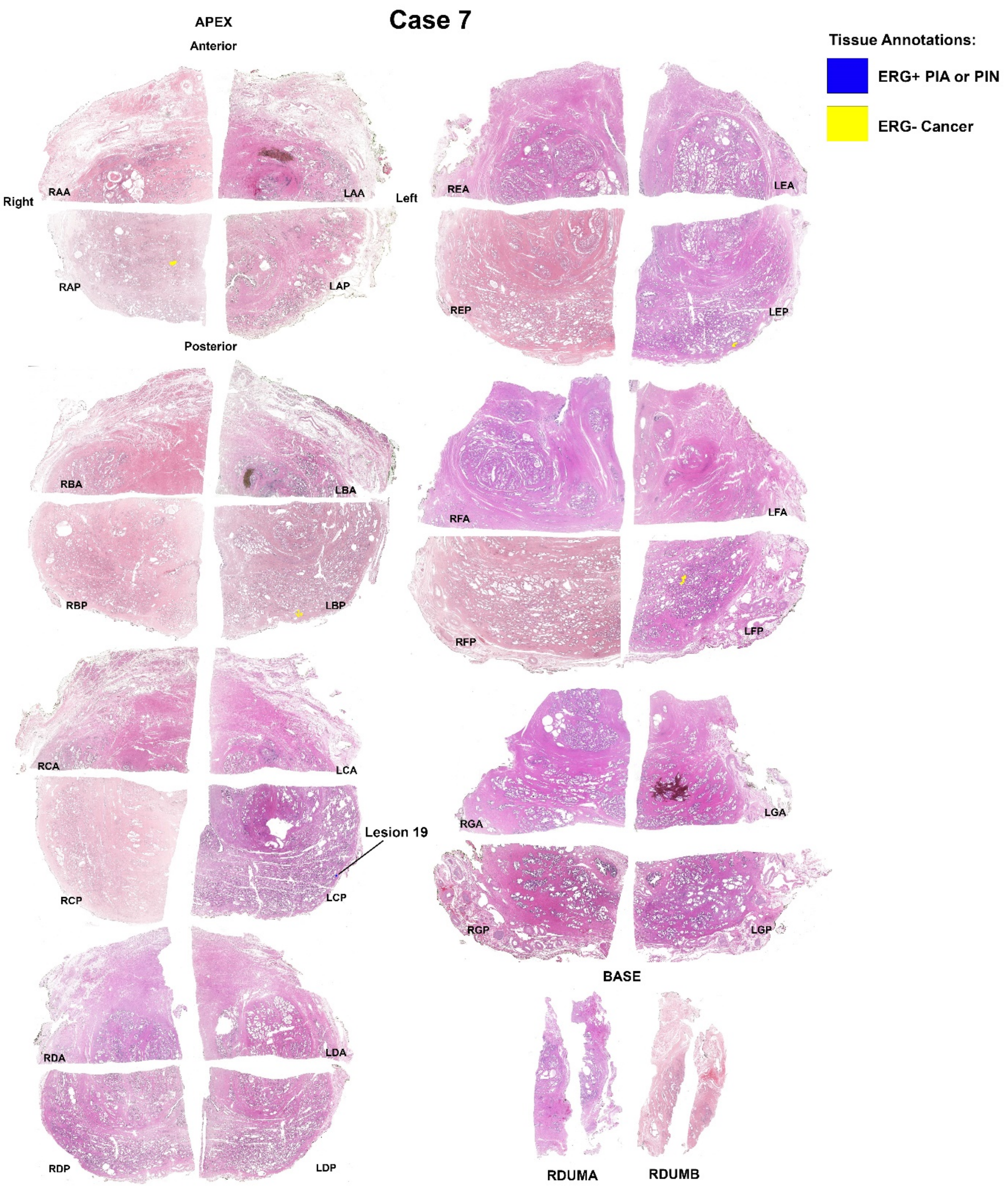


**Extended Data Fig. 13**. Prostate map showing location of ERG+ LGPIN (lesion 19) foci (blue) and ERG- cancer (yellow) in case 7. There was no ERG+ cancer found in this case.

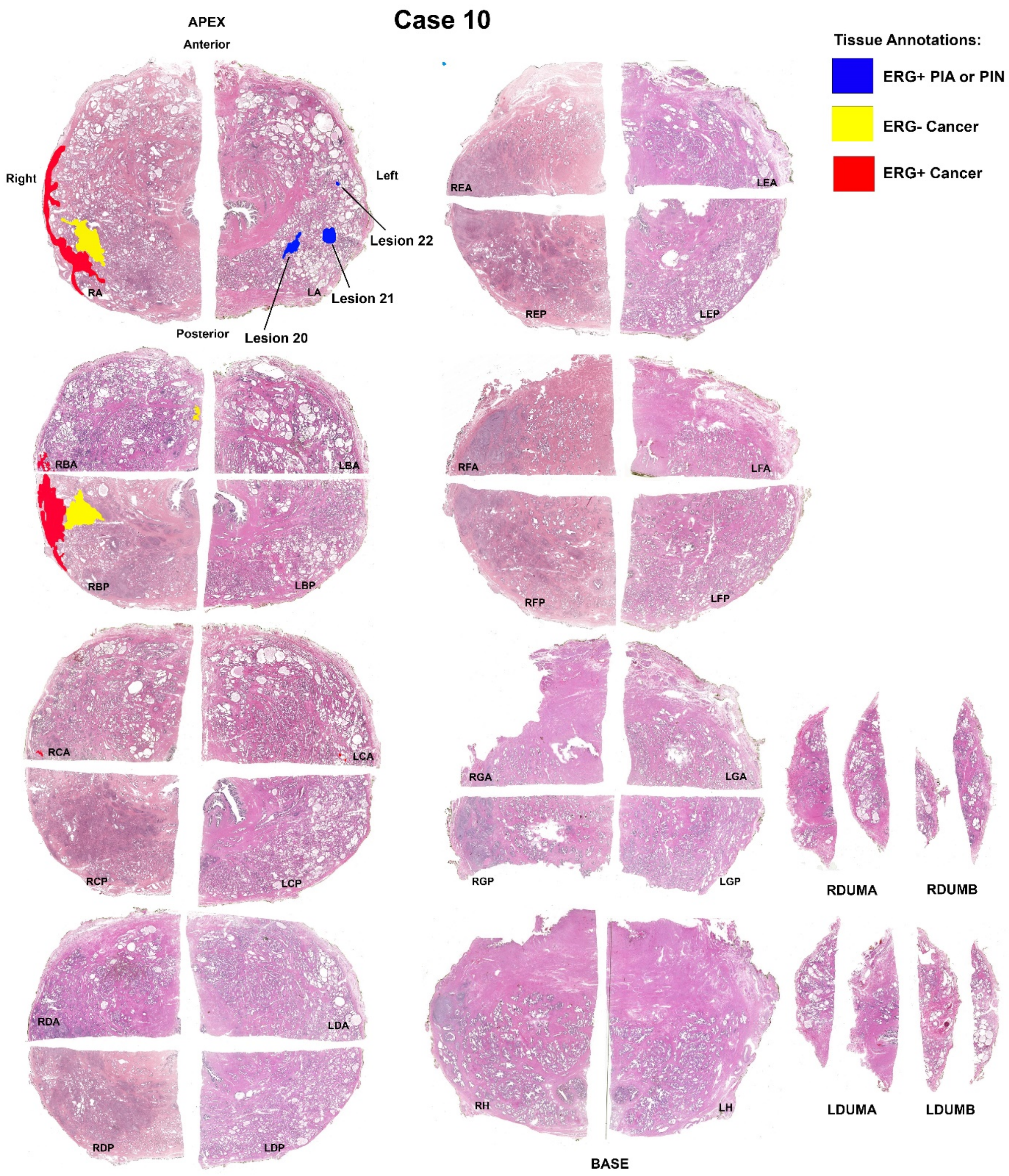


**Extended Data Fig. 14**. Prostate map showing location of ERG+ LGPIN/HGPIN (lesion 20 and 21), and PIA (lesion 22) foci (blue), ERG- cancer (yellow), and ERG+ cancer (red) in case 10.

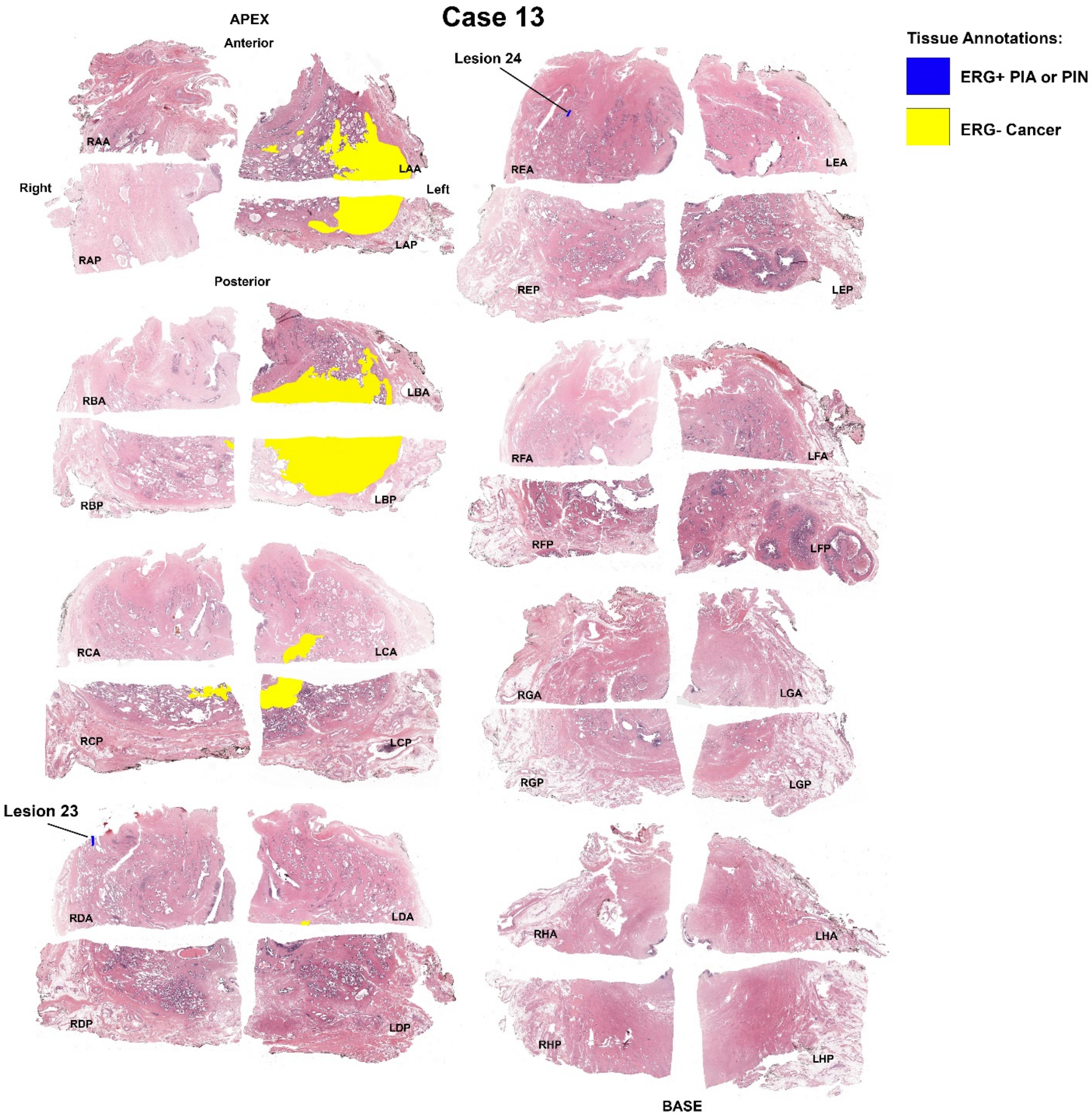


**Extended Data Fig. 15**. Prostate map showing location of ERG+ PIA (lesion 23 and 24) foci (blue) and ERG- cancer (yellow) in case 13. There was no ERG+ cancer found in this case.

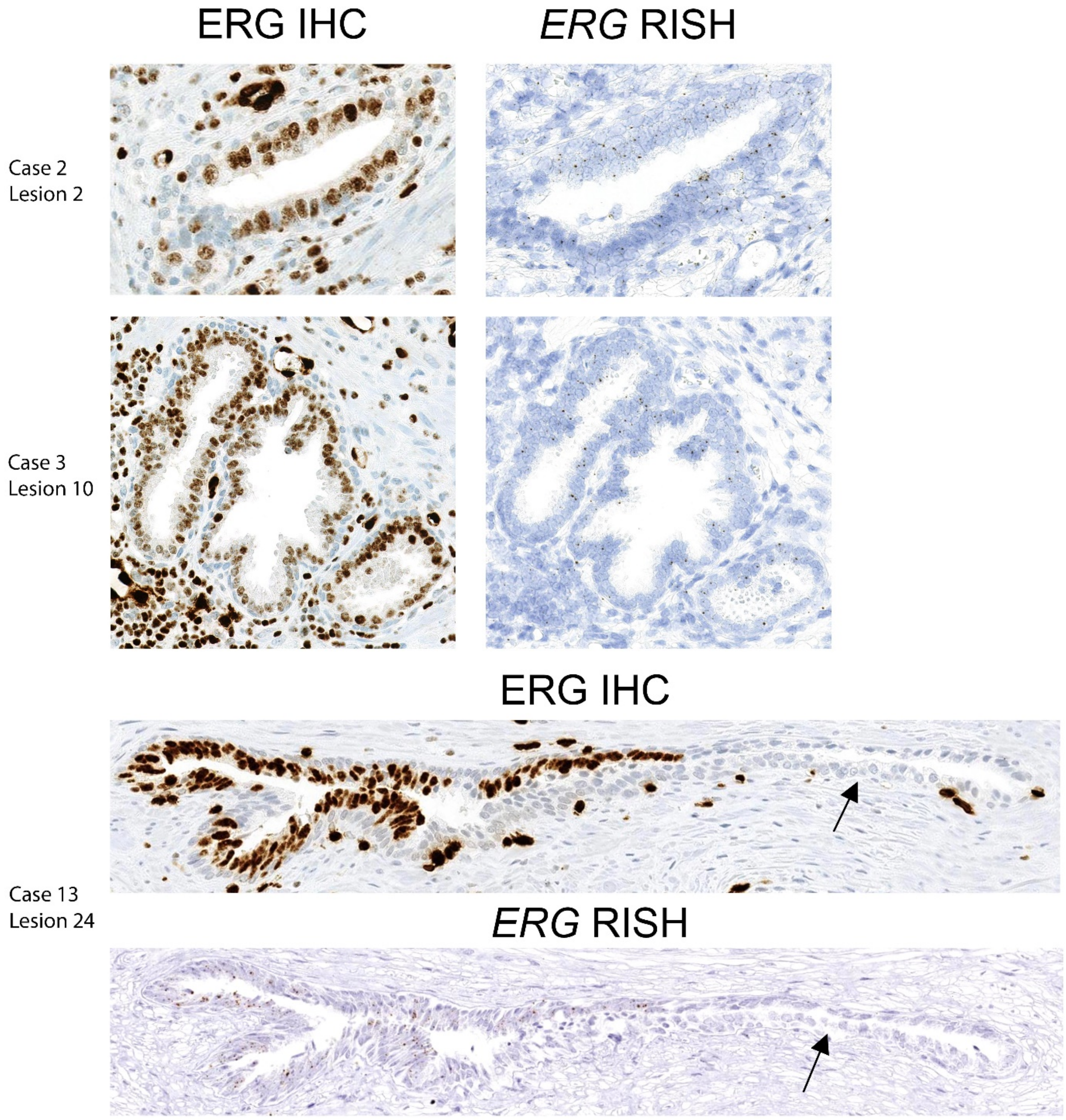


**Extended Data Fig. 16**. Additional examples demonstrating that PIA (lesion 2 and 24) and LGPIN (lesion 10) foci that are ERG+ by IHC are also positive for *ERG* mRNA by RISH. Arrows denote ERG- areas that are negative by both IHC and RISH.

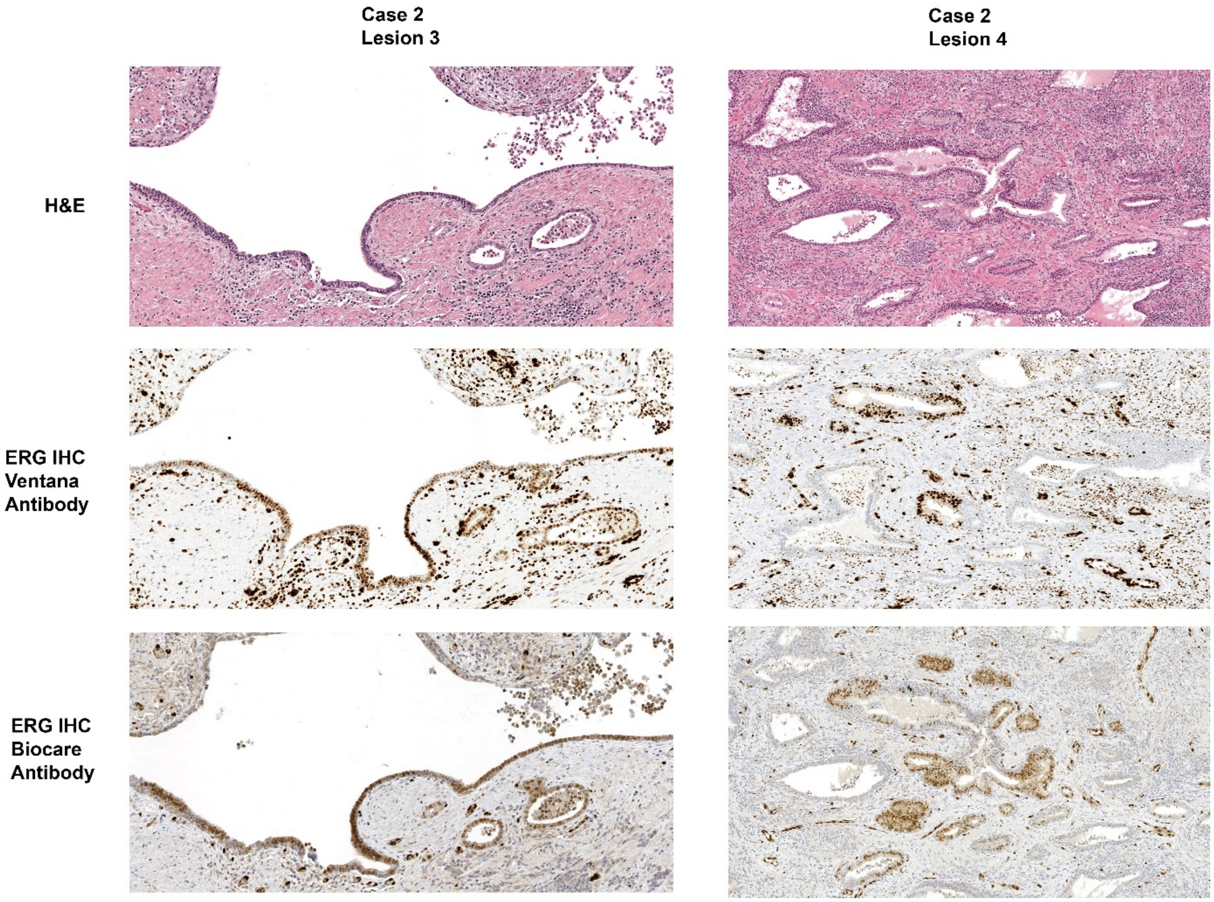


**Extended Data Fig. 17**. PIA foci that are ERG+ by IHC with the Ventana antibody (directed against C-terminus of ERG) are positive with a separate Biocare antibody (directed against N-terminus of ERG).

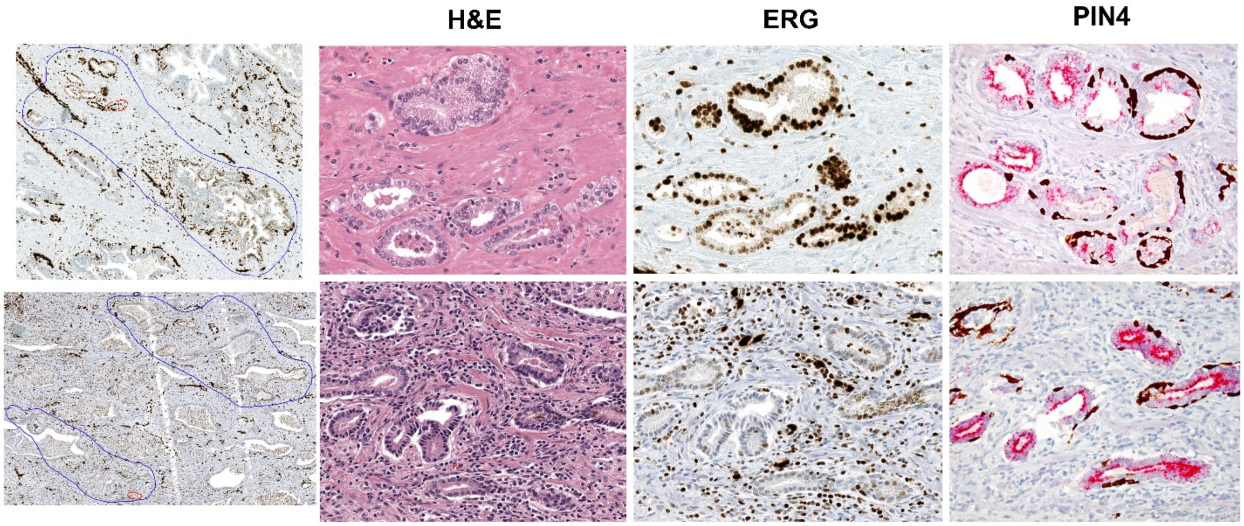


**Extended Data Fig. 18**. Additional examples of ERG+ microadenocarcinoma, apparently in the process of initial invasion, adjacent to ERG+ PIA. Large regions of ERG+ PIA (circled in blue in first panel of ERG IHC, 40X magnification) with small regions of adjacent microadenocarcinoma (circled in red) from case 3 (top row) and case 2 (bottom row). H&E, ERG IHC, and PIN4 IHC (all 200X magnification) demonstrate ERG+ PIA next to ERG+ microadenocarcinoma.

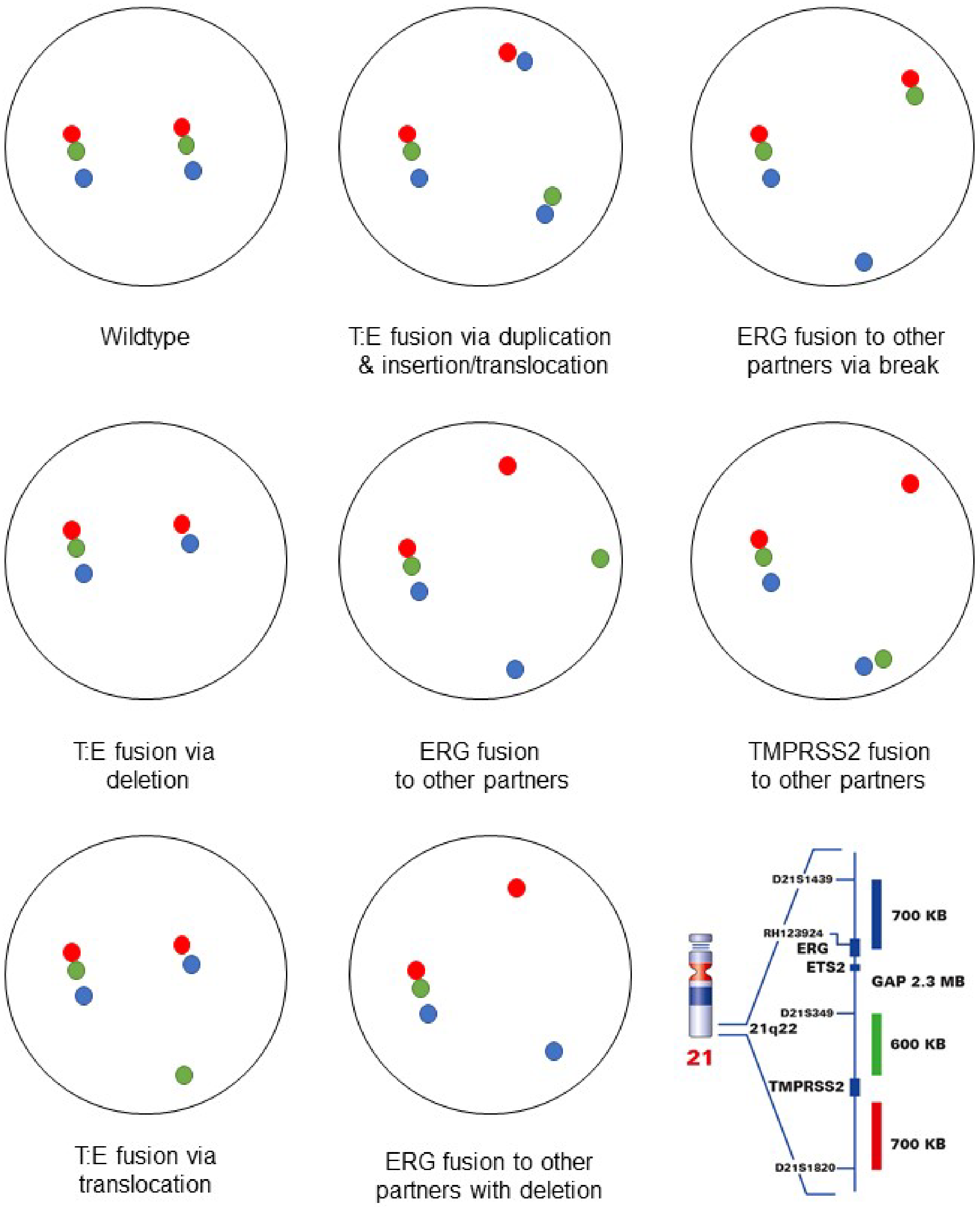


**Extended Data Fig. 19**. Examples of different fusion patterns that could be detected with the TMPRSS2:ERG (T:E) FISH assay. Red probe is located in the distal TMPRSS2 gene region, green probe is located in the proximal TMPRSS2 gene region, and blue probe is located in the ERG (21q22) gene region. Schematic of probe locations is from the Leica product information sheet (https://www.leicabiosystems.com/ihc-ish-fish/kreatech-fish-probes/on-probes/products/tmprss2-erg-21q22-ruo/)

**Extended Data Table 1**. Clinical and pathologic parameters of the infected radical prostatectomy cohort.

| **Case #** | **Age** | **Race*** | **Gleason** | **pStage** | **16S CISH** | **LPS IHC** | **LTA IHC** | **AFB** | **GMS** | **PKS (*clbB*)** | **Pathologic findings**** |
|---|---|---|---|---|---|---|---|---|---|---|---|
| 1 | 63 | W | 3+4=7 | T3BN0MX | Neg | Neg | Neg | N/A | N/A | N/A | Marked acute inflammation |
| 2 | 67 | W | 3+3=6 | T2NXMX | Pos | Pos | Neg | N/A | N/A | Pos | Florid non-specific granulomatous prostatitis with microabscess formation |
| 3 | 59 | W | 3+5=8 | T2N0MX | Pos | Pos | Neg | N/A | N/A | N/A | Acute inflammation and non-specific granulomatous prostatitis |
| 4 | 64 | W | 4+4=8 | T3AN0MX | Pos | Pos | Neg | N/A | N/A | N/A | Non-specific granulomatous prostatitis |
| 5 | 55 | W | 3+4=7 | T2N0MX | Neg | Neg | Neg | Neg | Neg | N/A | Marked chronic inflammation and granulomatous inflammation with small foci of necrosis |
| 6 | 66 | W | 4+3=7 | T3BN0MX | Neg | Neg | Neg | N/A | N/A | N/A | Marked acute inflammation |
| 7 | 57 | W | 3+3=6 | T2N0MX | Neg | Neg | Neg | N/A | N/A | N/A | Marked acute and chronic inflammation |
| 8 | 57 | O | 3+4=7 | T2N0MX | Pos | Pos | Neg | N/A | N/A | N/A | Extensive acute and chronic inflammation, inflamed urethra |
| 9 | 60 | W | 3+4=7 | T2N0MX | Neg | Neg | Neg | Neg | Neg | N/A | Non-necrotizing granulomas in prostate |
| 10 | 51 | B | 3+3=6 | T2NXMX | Pos | Neg | Pos | N/A | N/A | N/A | Granulomatous chronic inflammation in association with duct rupture |
| 11 | 52 | W | 3+4=7 | T2N0MX | Neg | Neg | Neg | Neg | Neg | N/A | Moderate to extensive non-caseating granulomata and acute and chronic inflammation |
| 12 | 65 | B | 3+4=7 | T2N0MX | Pos | Pos | Neg | N/A | N/A | Pos | Florid nonspecific granulomatous prostatitis |
| 13 | 67 | B | 3+4=7 | T2N0MX | Neg | Neg | Neg | N/A | N/A | N/A | Squamous metaplasia and cystitis cystica et glandularis involving the prostatic urethra with associated chronic inflammation |
| 14 | 57 | B | 3+4=7 | T2NXMX | Pos | Pos | Neg | N/A | N/A | N/A | Marked acute and chronic inflammation |
| 15 | 48 | W | 3+3=6 | T2N0MX | Neg | Neg | Neg | N/A | N/A | N/A | Granulomatous prostatitis |

*B = self-identified as Black or African American, W = self-identified as White or Caucasian, O = Other not self-identified as Black, White, American Indian or Alaska Native, Asian, or Native Hawaiian or Pacific Islander

Pos = positive, Neg = negative, N/A = not assessed, AFB = auramine/rhodamine stain, GMS = methenamine silver stain

**Pertains to uninvolved (non-tumor) areas of prostate, findings from diagnostic surgical pathology report

**Extended Data Table 2**. Pathologic assessment of all ERG+ PIA, LGPIN, and HGPIN foci.

| Case # | Block designation | Erg+ B9 lesion # | Notes |
|---|---|---|---|
| 2 | RAP | 1 | PIA, some reactive nuclei with nucleolar enlargement not diagnostic of LGPIN or HGPIN |
| | LAP | 2 | PIA, atypia*, some early invasive carcinoma (microadenocarcinoma**) |
| | RBP | 3 | PIA with nuclear reactive changes and microadenocarcinoma |
| | | 4 | PIA with nuclear reactive changes |
| | RCP | 5 | PIA |
| | RDP | 6 | PIA and microadenocarcinoma |
| | | 7 | PIA |
| | LDUMB | 8 | PIA with nuclear reactive changes and microadenocarcinoma |
| 3 | RAP | 9 | PIA |
| | RBP | 10 | LGPIN |
| | RCP | 11 | LGPIN |
| | | 12 | LGPIN |
| | | 13 | LGPIN |
| | | 14 | LGPIN, atypia, some glands microadenocarcinoma |
| | LCP | 15 | HGPIN |
| 4 | LDA | 16 | Merging PIA, LGPIN, and HGPIN. Atypia and microadenocarcinoma. |
| 5 | RDA | 17 | PIA |
| | RE | 18 | PIA and microadenocarcinoma |
| 7 | LCP | 19 | LGPIN |
| 10 | LA | 20 | LGPIN and HGPIN |
| | | 21 | LGPIN and HGPIN |
| | | 22 | PIA merging with atypia |
| 13 | RDA | 23 | PIA with nuclear atypia not diagnostic of LGPIN or HGPIN |
| | REA | 24 | PIA |

*atypia defined as small foci of atypical glands suspicious for but not diagnostic of carcinoma.
**microadenocarcinoma as defined by McNeal (*24*).
